# SEQUIN multiscale imaging of mammalian central synapses reveals loss of synaptic microconnectivity resulting from diffuse traumatic brain injury

**DOI:** 10.1101/704916

**Authors:** Andrew D. Sauerbeck, Mihika Gangolli, Sydney J. Reitz, Maverick H. Salyards, Samuel H. Kim, Christopher Hemingway, Tejaswi Makkapati, Martin Kerschensteiner, David L. Brody, Terrance T. Kummer

## Abstract

The complex microconnectivity of the mammalian brain underlies its computational abilities, and its vulnerability to injury and disease. It has been challenging to illuminate the features of this synaptic network due in part to the small size and exceptionally dense packing of its elements. Here we describe a rapid and accessible super-resolution imaging and image analysis workflow—SEQUIN—that identifies, quantifies, and characterizes central synapses in animal models and in humans, enabling automated volumetric imaging of mesoscale synaptic networks without the laborious production of large histological arrays. Using SEQUIN, we identify delayed cortical synapse loss resulting from diffuse traumatic brain injury. Similar synapse loss is observed in an Alzheimer disease model, where SEQUIN mesoscale mapping of excitatory synapses across the hippocampus identifies region-specific synaptic vulnerability to neurodegeneration. These results establish a novel, easily implemented and robust nano-to-mesoscale synapse quantification and molecular characterization method. They furthermore identify a mechanistic link—synaptopathy—between Alzheimer neurodegeneration and its best-established epigenetic risk factor, brain trauma.

## Introduction

Mammalian central neural networks are complex systems in which nanoscopic synaptic specializations precisely connect mesoscopic local circuit elements enabling the bottom-up emergence of behavior^1^. Understanding the cellular and circuit-level generation of such behavior therefore requires a detailed knowledge of synaptic architectures and phenotypes across multiple scales. Synaptic networks are also frequent targets of neurological injury, with pathological alterations in synaptic connectivity (synaptopathy) recognized as a central shared mechanism of neurodegenerative disease^2^. The clearest and most pressing example of this is the rising public health crisis of Alzheimer disease (AD)^3^. Extensive (25-40%^4,5^) and widespread synapse loss occurs early in AD^6,7^, and is better correlated with cognitive decline than classic AD neuropathologies such as amyloid plaques and neurofibrillary tangles^6,8^. Traumatic brain injury (TBI) is the best-established environmental risk factor for AD^9,10^, but its subcellular foci of injury and their connections to AD remain in question. Addressing these crucial and manifold challenges requires approaches to synaptic network imaging and analysis that are broadly-accessible, robust and quantitative, and extend the analysis of nanoscopic synaptic elements to mesoscale regions (generally millimeter scale networks that typify connectivity across individuals^11^).

Because of their size, dense packing, and the exceptionally complex subcellular environment in which they reside, synaptic structural analysis has been extremely challenging. Electron microscopic approaches have long been capable of identifying and quantifying individual synapses with nanometer resolution and excellent specificity informed by ultrastructural hallmarks (*e.g.*, postsynaptic densities and presynaptic vesicle clusters). Advanced implementations involving tape-collecting lathe ultramicrotomes^12^, serial block-face scanning^13^, or focused ion beam milling^14^ are capable of mapping synaptic connectivity across mesoscale circuits with exquisite precision. These approaches, however, suffer from severe limitations in speed, laboriousness, and availability: scan times for local circuits (*e.g.*, a murine cortical column) can easily extend to months or years^15^. Furthermore, the resulting voluminous imaging data must be annotated, frequently by hand, to catalogue structures of interest. Limited molecular phenotyping is a further challenge, with immunoelectron microscopy approaches largely limited to two-dimensional analysis and minimal multiplexing of markers.

In recent years array tomography (AT) emerged as a powerful extension of ultrasectioning-based synaptic characterization^16^. AT layers multiplexed immunofluorescent phenotyping onto electron microscopic ultrastructural analysis. Unlike standard optical sectioning techniques such as confocal microscopy that suffer from inadequate axial resolution to reliably separate neighboring synaptic elements, AT utilizes physical ultrathin sectioning (50-200 nm) to achieve axial super-resolution (S-R). AT has revealed numerous microstructural and molecular features of synapses^17^ and other nanoscopic structures^18^. However, its reliance on laborious and technically challenging production, staining, imaging, and subsequent reassembly of ultrathin two-dimensional tissue arrays remains a substantial impediment to widespread adoption^19^.

Optical implementations of S-R microscopy provide, in theory, a natively three-dimensional alternative. Existing approaches, however, are technically challenging in their own right, often require specialized fluorophores incompatible with transgenic approaches, and struggle with large or three-dimensional volumes, slow scan speeds, and/or weak signal strength^20^. Confocal laser scanning microscopy is diffraction limited but benefits from straightforward sample preparation, compatibility with conventional exogenous and endogenous fluorescent probes, and rapid imaging of large 3D volumes. A recently optimized innovation in sub-diffraction limited imaging, image scanning microscopy (ISM)^21,22^, combines lateral and axial sub-diffraction limit (*i.e.*, S-R) imaging with the well-developed labeling, hardware, accessibility, sensitivity, resistance to artifacts^20^, and speed of confocal microscopy. In the Airyscan implementation, this is accomplished via a novel 32-channel gallium arsenide phosphide photomultiplier tube (GaAsP-PMT) compound eye detector that reduces the effective confocal pinhole diameter while enhancing overall signal strength through parallelization of photon detection (Fig. 1A). Combined with appropriate image analysis workflows, Airyscan microscopy potentially represents an ideal balance of resolution, efficiency, and accessibility for the rapid assessment of synaptic networks at-scale.

**Fig. 1:**
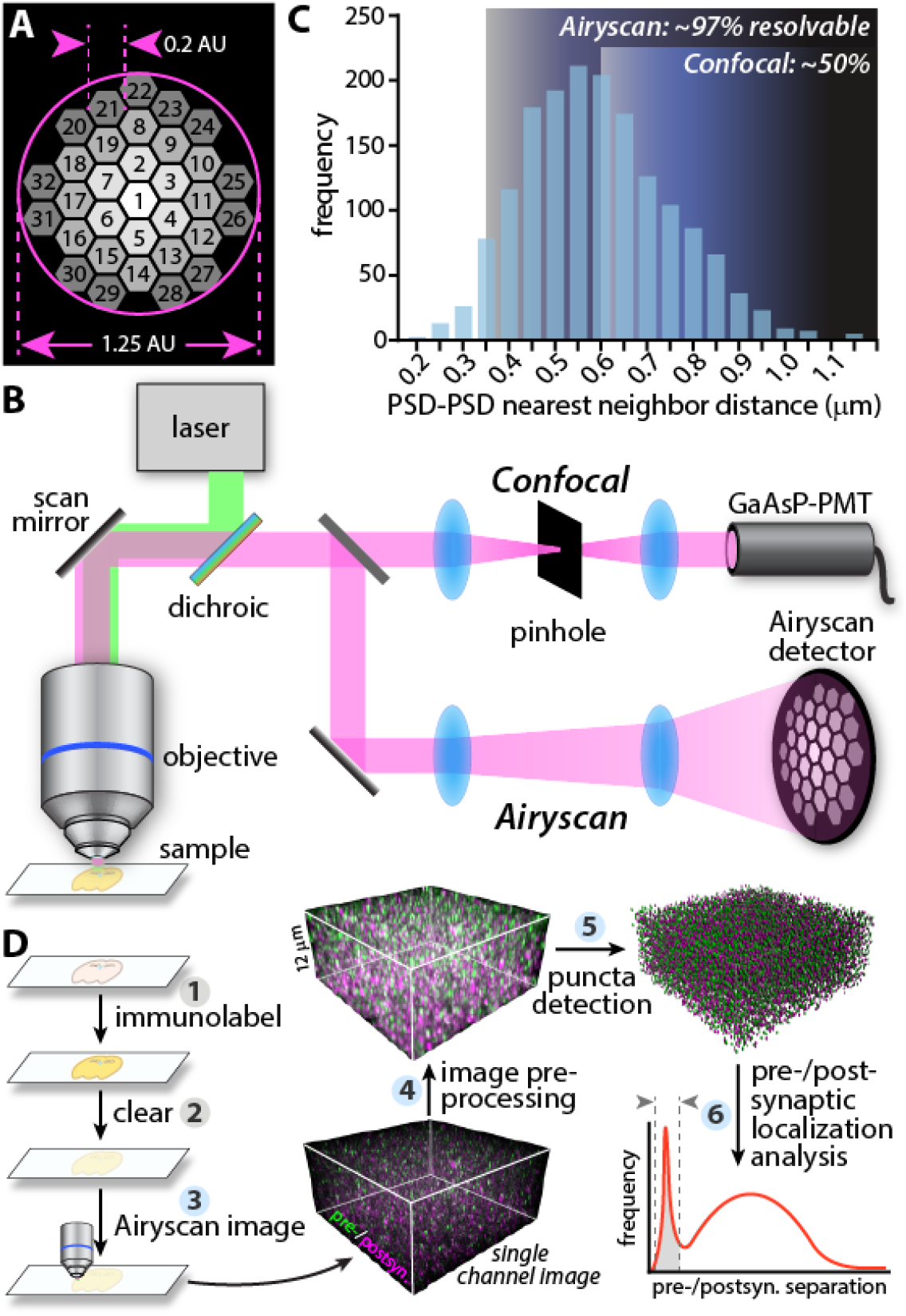
SEQUIN overview. (A) Schematic of compound eye Airyscan detector consisting of 32 unique detector elements each operating as a 0.2 AU pinhole. (B) Comparison of confocal and Airyscan light paths. (C) Inter-PSD separation by EM (secondary analysis of^12^) and resolving limit of confocal compared to Airyscan microscopes relative to these separations. Percent synapses resolvable in each case calculated from least resolved axis (axial, Z). (D) Overview of SEQUIN multiscale analysis method. Steps highlighted in blue are easily automated. Abbreviations: postsyn., postsynaptic.

We set out to develop new histological and image analysis methodologies to rapidly detect and quantify synaptic loci in mammalian neuropil. Our goal was to produce an easily implemented method that permitted quantitative analysis of nanoscale synaptic endpoints over mesoscale networks. The resulting ‘SEQUIN’ technique (Synaptic Evaluation and QUantification by Imaging of Nanostructure) is a combination of sub-diffraction limit, airyscan imaging and localization-based analysis that pinpoints pre- and postsynaptic elements and identifies synaptic loci by virtue of trans-synaptic separations in-line with ultrastructurally validated parameters. We demonstrate the utility of SEQUIN by analyzing synapse loss in an AD model and after diffuse TBI. By substantially lowering thresholds to perform large-scale synaptic quantification and derive population level feature sets, we expect that SEQUIN will broadly enable the application of synaptic connectivity and molecular phenotyping to critical questions in CNS health and disease.

## Results

### SEQUIN volumetric imaging of synaptic puncta in mammalian neuropil

Image scanning microscopy as implemented with the Airyscan microscope yields a 1.7-fold enhancement of lateral (XY) and axial (Z) resolution over standard confocal imaging^23,24^, improving resolution to 140 nm × 140 nm × 350 nm (XYZ). Super resolution is achieved by reducing the effective pinhole diameter well below 1 Airy unit (AU), while avoiding the dramatic loss of signal this would generally entail. The resulting light path lacks a physical pinhole aperture (Fig. 1A, B). Instead, each element of the compound eye detector acts as an independent 0.2 AU pinhole enhancing resolution, while post-imaging pixel reassignment ties sensitivity to the complete 1.25 AU array of detectors (Fig. 1A).

Ultrastructural analysis of mammalian cortex reveals an average separation of 595 nm between neighboring synapses^12^ (Fig 1C). This is below the axial resolution of diffraction-limited approaches such as confocal microscopy, resulting in poor identification of individual synaptic elements. A 1.7-fold enhancement in resolution, fortunately, is sufficient to resolve >97% of synaptic elements (*i.e.*, most synapses in mammalian gray matter are just below the axial resolution of diffraction-limited microscopy, Fig. 1C). Thus the robust improvements in optical sectioning afforded by ISM approaches such as Airyscan permit volumetric imaging of structures packed at synaptic density without resorting to laborious physical ultrathin sectioning.

The SEQUIN technique consists of (1) immunofluorescent labeling, (2) tissue clearing, (3) ISM of pre- and postsynaptic markers, (4) pre-processing of resulting 32-channel imaging data, (5) localization of pre- and postsynaptic centroids, and (6) identification of synaptic loci by virtue of ultrastructurally-informed separation distances (Fig. 1D). We initially chose to target synapsin and PDS-95—proteins that are abundantly expressed at central synapses where they play distinct roles in sculpting synaptic function^25,26^—as pre- and postsynaptic markers, respectively (Table S1). Imaging in thick sections revealed excellent antibody penetration to the center of 150 μm-thick sections (Fig. S1A, B).

Light scattering and refraction remain pervasive challenges in volumetric imaging, particularly with sensitive S-R applications^27,28^. SEQUIN was initially tested in 50 μm-thick murine brain sections without tissue clearing, but substantial degradation in image quality within a few microns of the tissue surface was observed (Fig. S1C). Clearing via refractive index matching in a water-soluble polyvinyl hydrocolloid (Mowiol 4-88) dramatically improved sensitivity and resolution, permitting excellent preservation of image quality 50 μm below the tissue surface (Fig. S1C, D).

Following immunolabeling and tissue clearing, we Airyscan imaged pre- and postsynaptic markers using high numerical aperture, oil immersion objectives. Dual channel SEQUIN data was acquired rapidly: 5.5 min were required to acquire a 53 × 53 × 12.6 μm image (< 0.01 sec/μm^3^). Image preprocessing entailed pixel reassignment and Weiner filter deconvolution (‘Airyscan processing’; Fig 1D, step 4), followed by correction of chromatic aberration using a within-experiment tissue, antigen, and fluorophore control (see Methods).

### Detection of synaptic loci via localization analysis

Image scanning microscopy and image pre-processing of a small region of mouse cortex resulted in a visually apparent improvement in sensitivity and resolution over confocal microscopy (Fig. 2A, B). As expected, densely-packed and less intense puncta were more readily distinguished using Airyscan vs. confocal microscopy, particularly those with nearest neighbors separated primarily in the axial dimension (Fig. 2A-C).

**Fig. 2:**
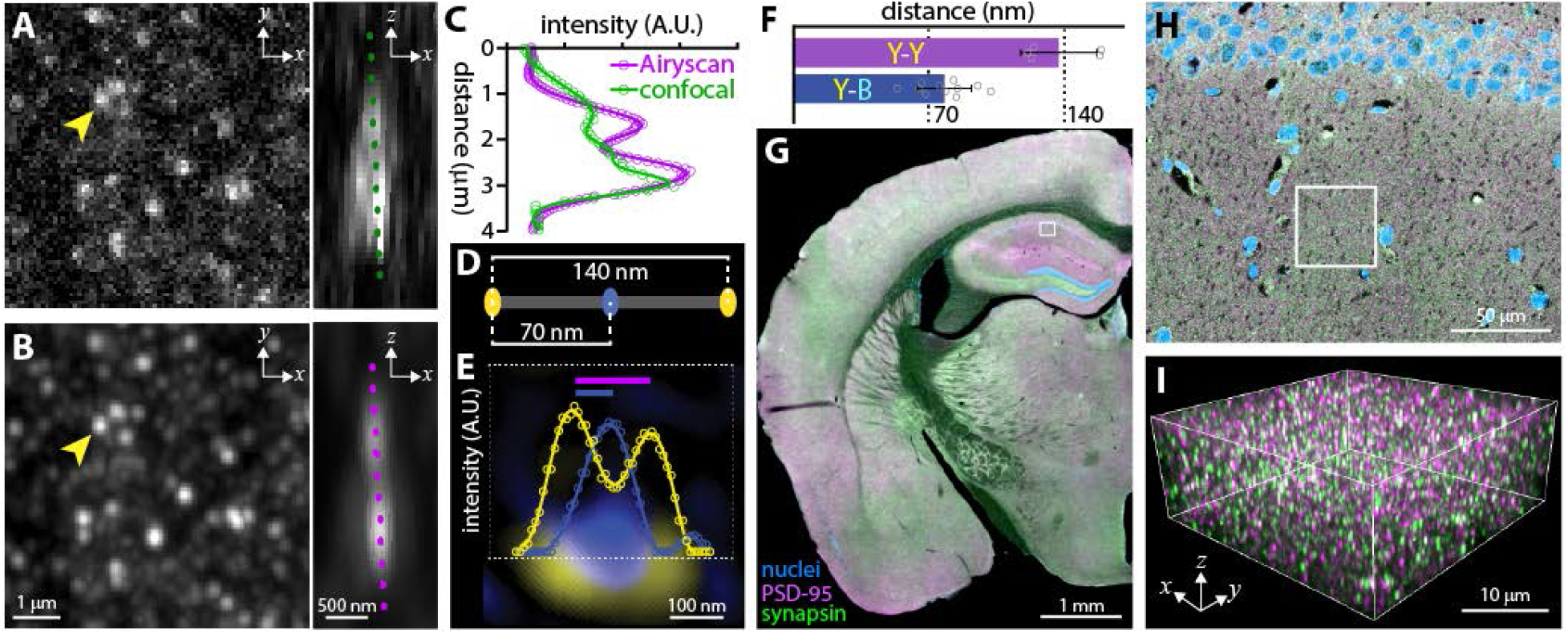
Resolution and localization analysis. (A) Maximally resolved confocal imaging of PSD-95 puncta. Arrowhead indicates punctum depicted to right with 90° rotation revealing a possible second punctum separated primarily in Z. (B) Same field imaged by ISM. Note improved resolution of puncta and enhanced sensitivity to dim puncta. Intensity profiles in (C) taken along dotted lines in (A) and (B) reveal improved separation of puncta in Z with ISM. (D) Schematic of DNA origami nanoruler. (E) Super-resolution image and intensity profile of nanoruler demonstrating relative localization of spectrally-distinct fluorophore centroids. Colored bars indicate measurements quantified in F. (F) Quantification of inter-fluorophore distances from E. Error bars +/−SEM from n = 5 nanorulers. (G) Low-power image of brain section labeled against pre- and postsynaptic markers with region in H indicated. (H) Higher magnification field from G; region depicted in I is indicated. (I) Boxed region from H with individual synaptic elements now visible. Fluorophore labels as per G.

In addition to resolving puncta from their neighbors, identifying synaptic loci by virtue of pre-to-postsynaptic marker separation requires localization of individual puncta accurately to within tens of nanometers^29^. Fortunately, localization microscopy approaches permit localization of spectrally-distinct fluorophores relative to one another at distances substantially less than the resolution limit of the microscope^30,31^. To specifically evaluate localization performance in the expected range of separation (10s-100s of nanometers^29^), DNA origami nanorulers with spectrally distinct fluorophores separated by 70 and 140 nm were imaged using ISM (Fig. 2D, E). The measured distance between these fluorophores agreed closely with their known physical separation (Fig. 2F), confirming the accuracy of SEQUIN localization measurements at distances in-line with pre-to-postsynaptic marker separations.

Our next goal was to replicate this localization analysis over all imaged pre- and postsynaptic puncta throughout a volume of murine cortex (Fig. 2G-I; see Fig. 1D, step 5). To accomplish this, we located the centroids of each pre- and postsynaptic punctum within the three-dimensional volume (Fig. 3A-C). Closer inspection of the resulting intensity-filtered point patterns using spatial statistics (Ripley’s K function^32^) revealed that pre- and postsynaptic puncta were distributed statistically indistinguishably from random (Fig. 3D, E). When the distribution of presynaptic puncta was considered relative to postsynaptic puncta, however, a clear tendency to co-associate emerged (Fig. 3F).

**Fig. 3:**
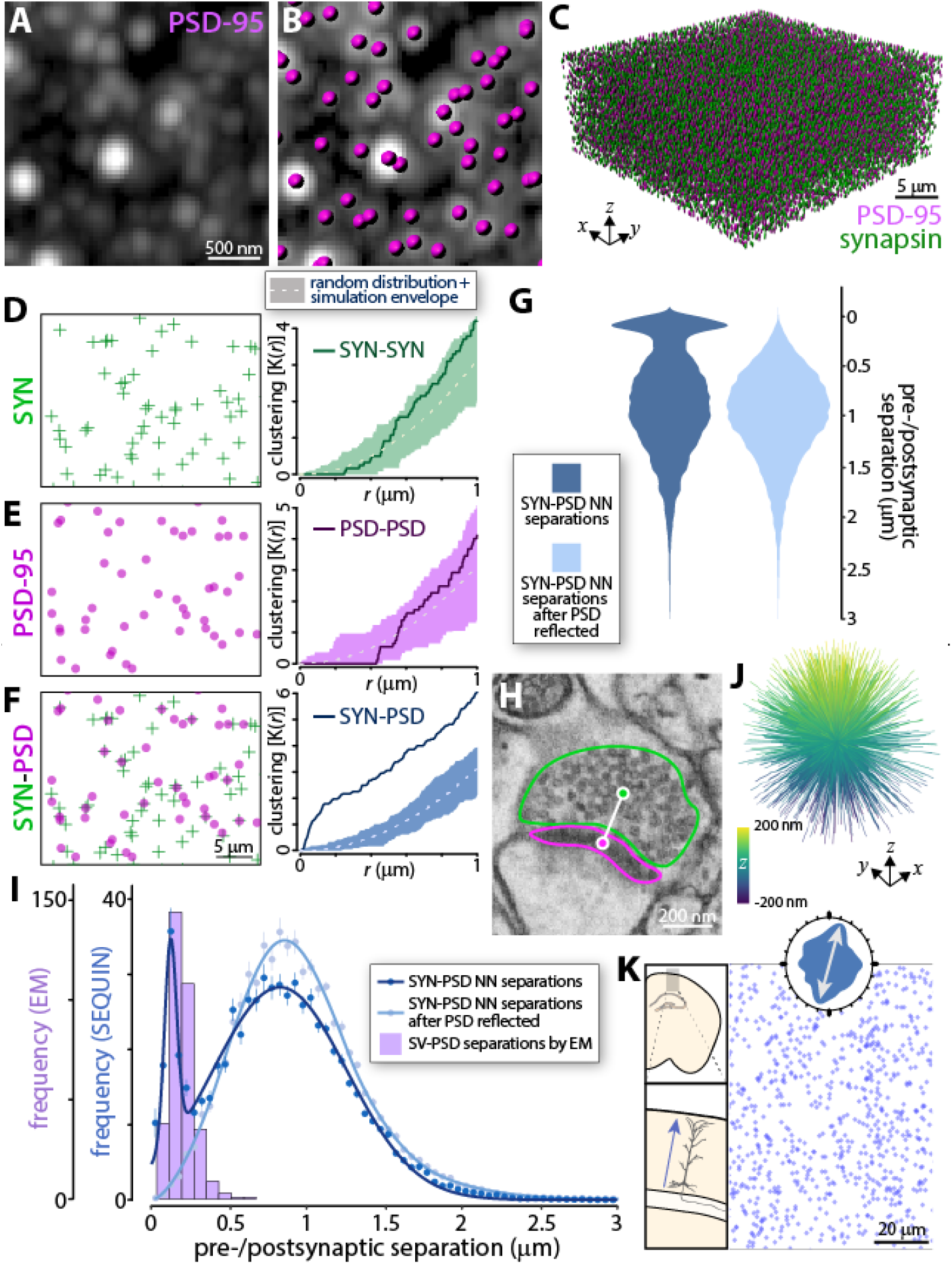
SEQUIN detection of synaptic loci. (A-B) Individual postsynaptic puncta centroids are located in 3-dimensions (magenta footballs) and added to presynaptic centroids (green footballs) to construct a volumetric map of synaptic elements in murine cortex (C). Spatial analysis of presynaptic (D; left panel) and postsynaptic (E; left panel) point patterns using Ripley’s K function (D, E; right panels). Within-marker spatial associations are indistinguishable from random. (F) Between-marker spatial analysis of pre- and postsynaptic point patterns demonstrates co-association above random. (G) Violin plots of separation distances between pre- and postsynaptic elements by SEQUIN localization analysis reveals a bimodal distribution of separations, which is lost when the postsynaptic point pattern is reflected about the Y axis. (H) Example of electron microscopic image used to measure distance (white line) between PSD (magenta outline) and synaptic vesicle reserve pool (green outline). (I) Frequency distribution of SEQUIN pre-to-postsynaptic separations before and after reflection of the postsynaptic point pattern compared to that measured by EM. Note correspondence between distances measured by EM and the early peak of the SEQUIN analysis. Error bars +/−SEM. (J) Intra-synaptic vector orientations across a volume of murine cortex lacks orientation bias, consistent with expected anatomy. (K) Synaptic loci distribution in layer II/III murine cortex. Inset, rose plot of nearest neighbor orientations (inter-synaptic orientation distribution function) reveals tendency for synaptic loci to be located along radial trajectories consistent with anatomy of pyramidal dendrites (see schematic to left). Examples representative of n ≥ 3 animals experiments for all panels. Abbreviations: SYN, synapsin; NN, nearest neighbor; SV, synaptic vesicle.

We therefore identified the nearest trans-synaptic partner (*e.g.*, the nearest neighbor of the opposite type) for each punctum, and calculated the Euclidean distance in three dimensions between these partners for the entire volume. This was accomplished using custom SEQUIN scripts written in MATLAB that accompany this report. Examination of these relationships revealed a bimodal distribution of separations (Fig. 3G), with an early concentration centered at approximately 150 nm pre-to-postsynaptic marker separation, and a broad concentration consisting of more distant associations. Suspecting that the early peak represented paired puncta at synaptic loci while the second peak consisted of random associations, we repeated the analysis after reflecting the PSD-95 image about its Y axis. This maneuver maintained puncta density and overall distribution while eliminating putative trans-synaptic relationships. As expected, the resulting distribution lacked the early, but retained the broad late peak of more distant associations (Fig. 3G), indicating that these distant associations are non-synaptic. The early peak, in contrast, represents non-random, nanoscopic associations of pre- and postsynaptic markers consistent with synaptic loci. Closer examination of the bivariate (synapsin-to-PSD-95, Fig. 3F) K function confirmed this interpretation: non-random associations appeared at short separation distances (<200 nm separation), but only random associations appeared at longer separations (the difference between K(*r*) for synapsin/PSD-95 and for a random distribution remains constant).

Prior analyses using STORM^29^ and STED^33^ indicate that pre- and postsynaptic markers associate at distances consistent with the early peak in the SEQUIN localization analysis. To further examine the nature of these closely separated pairings, we measured separations between post-synaptic densities and synaptic vesicle reserve pool centroids using electron microscopy. These ultrastructural features are expected to correspond to the locations of the PSD-95 and synapsin centroids, respectively, in the SEQUIN analysis (Fig. 3H)^34^. The distribution of inter-marker separations in this ultrastructural dataset (n = 339 synapses, median 196 nm, SD 79 nm) closely overlapped with the sharp, early peak in the SEQUIN analysis (Fig. 3I), further indicating that this peak represents synaptic loci. Importantly, the appearance of a bimodal distribution of pre-to-postsynaptic separations was not a result of spectral cross-talk, antibody-related or image processing artifacts, and was replicated using the same methods in an independent laboratory at a separate institution (Fig. S2).

Several additional features of closely-separated pre-to-postsynaptic pairs were consistent with synaptic loci. The intra-synaptic directional orientation (pre-to-postsynaptic centroid orientation vector) of such synaptic loci lacked bias (Fig. 3J; relative anisotropy of resulting orientation distribution matrix = 0.006), in-line with anatomic expectations and despite the non-isotropic resolution of ISM. Moreover, the inter-synaptic distribution of synaptic loci in murine cortex exhibited a tendency for radial anisotropy, consistent with the predominant orientation of pyramidal dendritic arbors perpendicular to the cortical surface (Fig. 3K and Fig. S3). Detection of synaptic loci using SEQUIN is furthermore robust to overall image intensity (Fig. S4), in contrast to methods dependent on image thresholding or densitometry. Thus SEQUIN is well suited for comparisons of tissues with differing labeling efficiency, background levels, or opacity (*e.g.*, Fig. 5).

### Quantification of synaptic density via SEQUIN analysis

To achieve the greatest labeling efficiency, antibodies against pre- and postsynaptic proteins were tested, yielding a cocktail of polyclonal pre- and postsynaptic synapsin and PSD-95 antibodies that maximized detection. Inspection of the distribution of pre-to-postsynaptic separation distances as a function of puncta intensity revealed that the majority of pairings consistent with synaptic loci were formed by relatively bright puncta (both pre- and postsynaptic; Fig. 4A, B)—dimmer puncta were more likely to pair at longer separations consistent with random associations. All puncta intensity groups, however, contributed puncta pairs separated at distances characteristic of synapses.

**Fig. 4:**
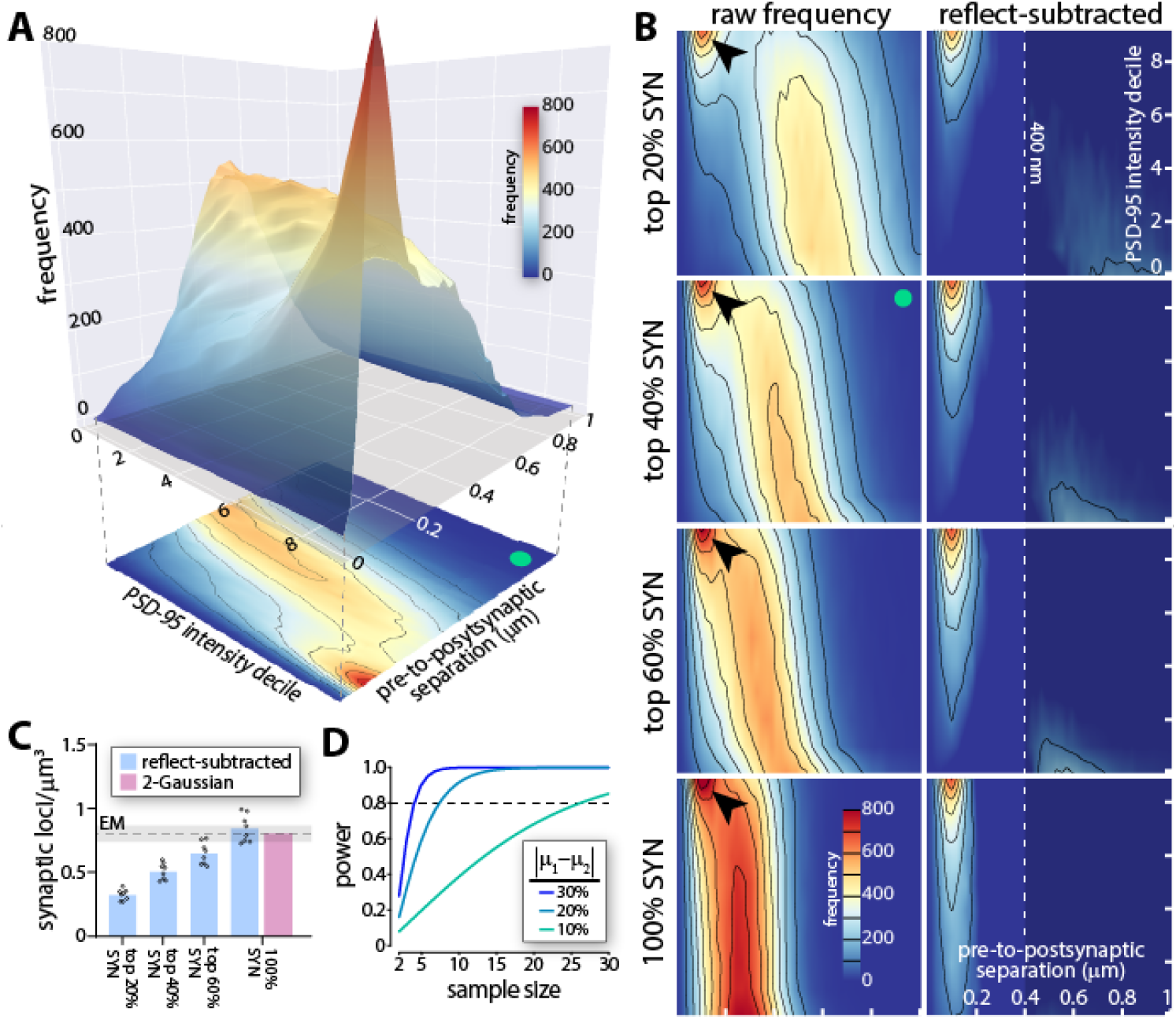
Synaptic density quantification. (A) Comparison of pre-to-postsynaptic separation distances as a function of intensity decile of PSD-95 puncta. Peak in separations consistent with synapses (red) is most pronounced with brighter PSD-95 puncta. 2D projection shown below exemplifies convention used to generate heatmaps in B (green dot, identical dataset in A, B). (B) Heatmaps of pre-to-postsynaptic separations vs. PSD-95 intensity decile with different synapsin puncta intensity bins. As additional synapsin puncta are added to the analysis, the more distantly separated random association peak migrates left (closer separations due to greater overall density of puncta). The early peak in associations consistent with synapses remains identifiable (arrowhead), but becomes challenging to separate from random associations in more inclusive datasets (*e.g.*, bottom left panel). Subtracting reflected frequency distributions that contain only random associations isolates pairings consistent with synapses (right panels). Dotted line (400 nm; right panels) used as cut-off for quantification in C. (C) Quantification of synaptic loci identified using increasingly inclusive synapsin puncta intensity bins after reflect subtraction or unmixing of Gaussian distributions. Mean and range of several EM estimates^12,31–33^ shown. n = 7 animals. (D) Power analysis of synaptic quantification in murine cortex (analysis for Ab cocktail shown). Abbreviations: SYN, synapsin.

Early and late peaks overlap substantially in such intensity-inclusive datasets (Fig. 4B). To quantify synaptic loci we therefore isolated anatomical colocalizations (those dependent on subcellular microstructure) from those formed randomly (those dependent only on puncta density and distribution) by subtracting the purely random colocalizations identified after reflection of the PSD-95 image about the Y axis (Fig. 4B, and see Fig. 3I). Quantification of resulting synaptic loci at separation distances ultrastructurally consistent with synapses (< 400 nm; see Fig. 3I) closely agreed with prior estimates from electron microscopy (0.85 vs. 0.82 synapses/μm^3^ EM^12,35–37^; Fig. 4C). Disentangling the SEQUIN puncta separation distribution through modeling as a mixture of two Gaussians generated a remarkably similar estimate (0.81 synapses/μm^3^; Fig. 4C and S6). Reassuringly, there was minimal difference when using a post- (*i.e.*, Fig. 4) or presynaptic marker as the reference dataset for pairing (0.847 synapses/μm^3^ PSD-95 reference vs. 0.854 synapses/μm^3^ synapsin reference, p = 0.90; Fig. S7). Variability in the SEQUIN analysis was modest (coefficient of variation 12.1-12.7%, pre-/postsynaptic reference pairing), resulting in practical sample size estimates for meaningful differences in synaptic density (n = 6.9 for a 20% difference at 80% power, alpha 0.05 for single antibody markers; n = 7.4 for the antibody marker cocktail; Fig. 4D). Although the antibody cocktail maximized detection of putative synaptic loci, antibody species multiplexing against PSD-95 limited flexibility for additional markers and required numerous additional controls for antibody cross reactivity (Fig. S5). Subsequent experiments were therefore conducted with single species primary antibody combinations.

### Pathological loss of synaptic loci measured with SEQUIN

To test the ability of SEQUIN to detect synaptopathic alterations, we turned to a murine model of Alzheimer disease (APPswe/PSEN1dE9^38,39^, or APP-PS1), a neurodegenerative condition characterized by diffuse and amyloid plaque-associated synapse loss^40,41^. Cortical synaptic density in control animals (C57Bl/6) was compared to that of aged (18 mo) APP-PS1 mice in the immediate vicinity of amyloid plaques and in cortex without respect to plaque location. As expected^42,43^, SEQUIN revealed a proximity-dependent loss of synapses in the vicinity of plaques (Fig. 5A, B). Interestingly, quantifying synaptic elements diffusely in cortex revealed only modest alterations in pre- or postsynaptic puncta compared to controls (Fig. 5C-E; presynaptic 1.3% decline, Cohen’s *d* = 0.12; postsynaptic 9.8% decline, Cohen’s *d* = 1.78), despite an overall reduction in signal intensity in the APP-PS1 animals compared to controls. The expected loss of synaptic loci, however, emerged with SEQUIN analysis (Fig. 5F; 41.8% decline; Cohen’s *d* = 2.75).

**Fig. 5:**
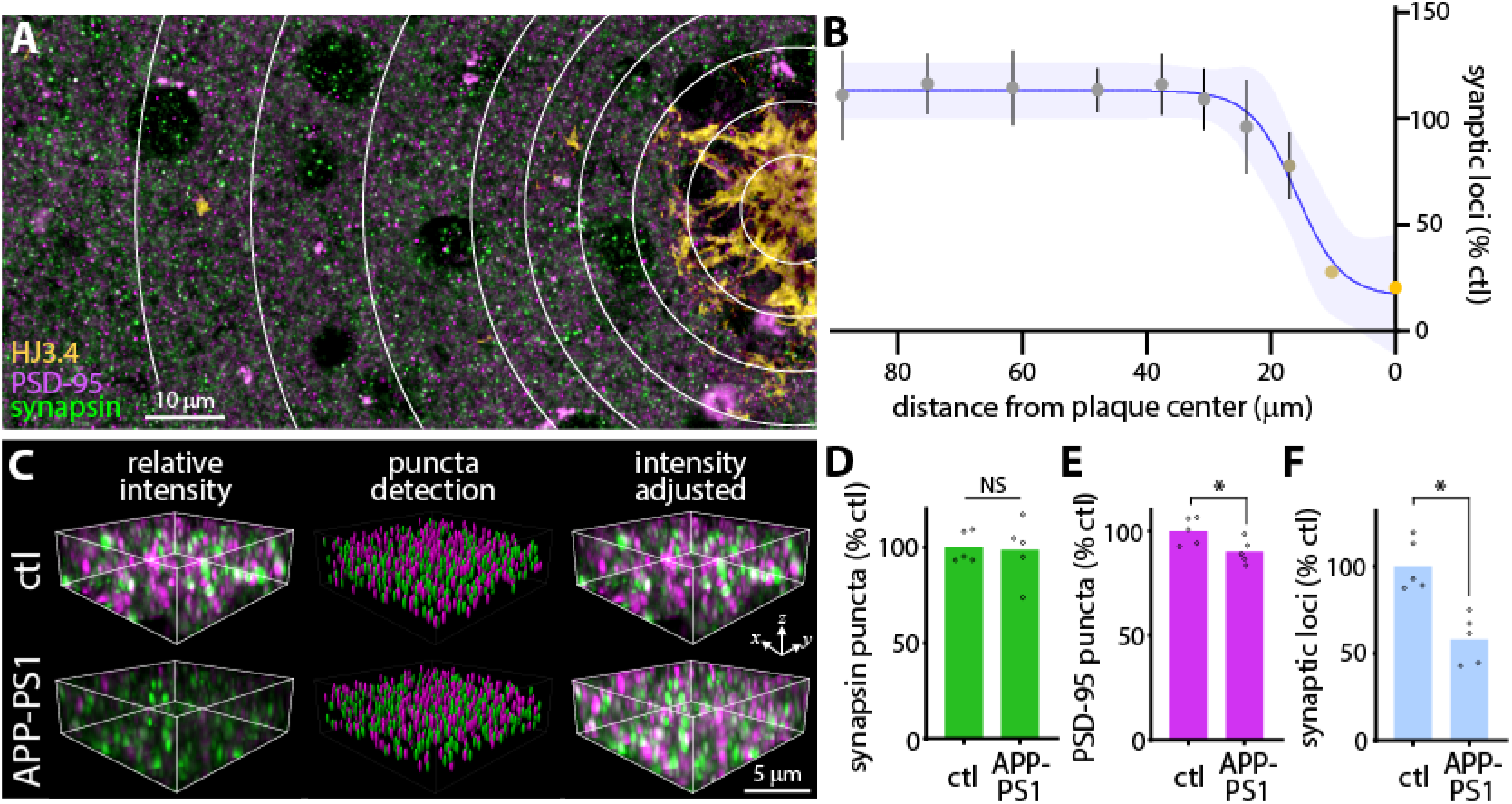
Synapse loss in an Alzheimer disease model. (A) Cortical region containing single amyloid plaque (HJ3.4 labeling). Concentric rings highlight regions used for quantification of synaptic loci in B. (B) Synaptic loci as a function of distance from plaque center with 95% CI of trend line indicated. Error bars +/−SEM from n = 7 plaques from 3 animals. (C) pre- and postsynaptic marker images from control and APP-PS1 cortex acquired with identical parameters (left panels) reveal less intense puncta in AD model, particularly postsynaptic puncta (magenta). Puncta detection (middle panels) is similar between groups, however, and intensity adjustment reveals a similar underlying pattern of puncta (right panels). Fluorophore labels as per A. (D, E) Quantification of puncta numbers from unadjusted images in C demonstrating similar numbers of puncta across conditions. (F) SEQUIN quantification of synaptic loci. n = 5 animals/group D-F. * p < 0.05. Abbreviations: ctl, control.

TBI is associated through unknown pathological mediators with a substantially and dose-dependent increased risk of developing a dementing illness, particularly AD^9,44,45^. To evaluate potential neuropathological connections between brain trauma and AD, we examined synaptic health after TBI using SEQUIN. TBI was induced using a tunable model of mild-moderate closed head diffuse TBI called modCHIMERA (Fig. 6A, B)^46^. Unlike predominant TBI models, modCHIMERA does not induce cortical neuronal loss, diffuse atrophy or cortical atrophy (Fig. 6C and S8), yet does result in neurobehavioral deficits indicative of neural circuit interruption^46^. Brains were collected and processed at intervals (1-30 days) post-injury (Fig. 6D, E). SEQUIN analysis revealed that synaptic loci were preserved at acute time points but were substantially reduced by 7 days post injury (Fig. 6F; 22.3% decline, Cohen’s *d* = 1.29). This effect persisted at least 1 month later (25.9% decline at 30 days, Cohen’s *d* = 1.60). To validate these results against gold standard (but far more laborious) methodology, electron microscopy was undertaken in a small parallel cohort. Synapses from animals subjected to modCHIMERA TBI exhibited several dystrophic features at 7 days post injury, including diffuse electron dense presynaptic material, enlarged synaptic vacuoles, and degenerating presynaptic boutons (Fig. 6G and S9), that have been observed in Alzheimer-related and other forms of neurodegeneration^47–49^. Stereological quantification of L1 cortical synapses by EM paralleled findings from SEQUIN quantification in magnitude (Fig. 6H; 27.1% decline, Cohen’s *d* = 2.37), though the analysis was underpowered. Thus SEQUIN provides insight into pathological alterations of synaptic microconnectivity across acute and chronic neuropathological conditions.

**Fig. 6:**
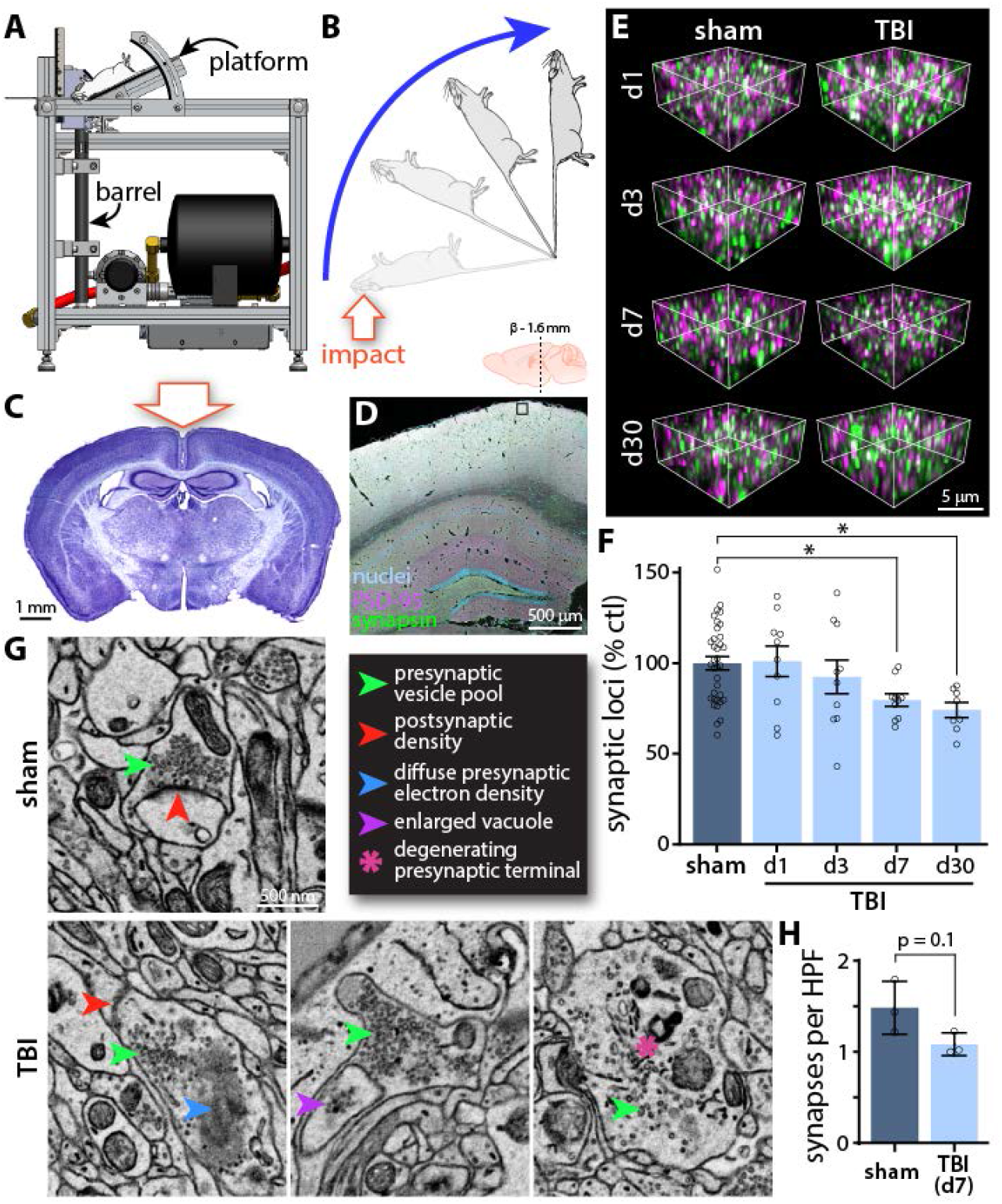
Cortical synapse loss after diffuse closed-head TBI. (A) Schematic of the CHIMERA impactor device^41,96^. Mouse rests on platform with head over barrel. Piston travels up barrel striking head of mouse. (B) Rotational movement of mouse resulting from impact. Mouse travels 180° coming to rest on a padded platform. (C) Appearance of brain 7 days after modCHIMERA diffuse TBI. Note lack of focal cortical pathology. (D) Overview of immunolabeling with location of coronal section and example of imaged region in layer 1 cortex boxed (images collected bilaterally). (E) Example imaged sub-volumes from regions near boxed area in D at various time points post-injury. Fluorophore labels as per D. (F) SEQUIN quantification reveals loss of synaptic loci by 7 days that persists to at least 30 days post-TBI compared to controls. n ≥ 8 animals/group. (G) Example of intact synapse in control animal by EM and examples of dystrophic synapses identified 7 days post modCHIMERA TBI. Scale bar applies to all panels. (H) Quantification of synapses by EM in control animals and after TBI. n = 3 animals/group. * p < 0.05. Abbreviations: HPF, high power field.

### Analysis of distinct synaptic subsets

Synapses are multicomponent machines whose functional properties are determined by their specific wiring patterns and molecular composition. A complete analysis of synaptic connectivity requires the means to capture such diversity. To evaluate this potential, SEQUIN was applied to subsets of synapses with alternate afferent origin (vGluT2+ thalamocortical afferents^50^) and physiological properties (inhibitory synapses). Examination of the pattern of vGluT2+ synaptic loci in murine cortex revealed a concentration in layer 4 consistent with known laminar specificity of thalamocortical axon termination in somatosensory cortex (Fig. 7A, B). Labeling against the inhibitory postsynaptic marker gephyrin along with the pan-presynaptic marker synapsin permitted quantification of inhibitory synaptic elements using SEQUIN (Fig. 7C). As demonstrated recently^51^, quantification of relative proportions of obligatory synaptic proteins can enable deeper analyses of synaptic networks, revealing anatomical and functional aspects of a network’s connectivity from the structural properties of its nodes (see Discussion). We therefore examined the alternate postsynaptic excitatory marker Homer 1 using SEQUIN. Similar to analysis involving PSD-95, pairing of synapsin with Homer 1 labeling enabled detection of excitatory synaptic loci (Fig. 7D).

**Fig. 7:**
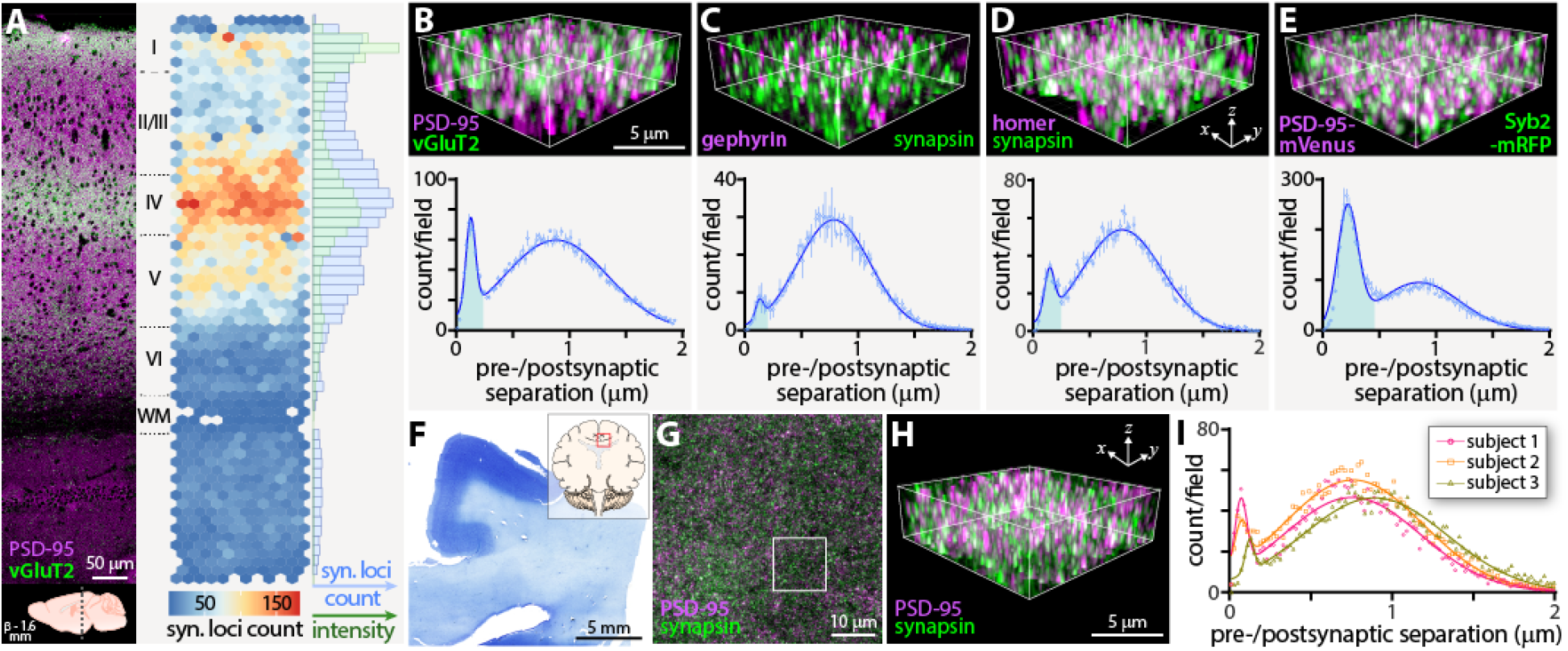
Application of SEQUIN to diverse synaptic populations. (A) Tile scan image of murine cortex labeled against vGluT2 and PSD-95 with lamina identified. Location of section relative to Bregma noted below. Adjacent heatmap indicates vGluT2+ synaptic loci count across cortical depth. Marginal histogram quantifies vGluT2+ synaptic loci (blue) and intensity of vGluT2+ axon terminals (green). (B) Example of quantified cortical subregion and frequency distribution of pre-to-postsynaptic puncta separations for vGluT2+ synaptic loci. Early peak consistent with synaptic ultrastructure highlighted in blue B-E. (C) Labeling of the inhibitory postsynaptic marker gephyrin and the pan-presynaptic marker synapsin and resulting quantification. (D) Quantification of synaptic loci using the alternate excitatory postsynaptic marker Homer 1. (E) Use of double knock-in mice for pre- and postsynaptic marker fluorescent fusion proteins (Syb2-mRFP and PSD-95-mVenus) to quantify synaptic loci. n = 3 animals each, A-E. (F) Low power image of human brain section used for quantification (subject 1) with location noted in inset. (G) Overview of labeled pre- and postsynaptic puncta, with region shown at higher magnification in H boxed. (H) High power image of synaptic elements in human brain for comparison with labeling in murine tissue. (I) Frequency distribution of pre-to-postsynaptic puncta separations for three human subjects demonstrating identification of early peaks consistent with synaptic ultrastructure.

Intravital analysis of synaptic alterations requires endogenous and/or vital labeling of synaptic structures. To evaluate the suitability of SEQUIN for such an application, we generated a double knock-in mouse for spectrally-distinct pre- and postsynaptic fluorescent fusion proteins (Syb2-mRFP^52^ and PSD-95-Venus^53^). SEQUIN imaging and analysis of tissue from these animals revealed punctate labeling of pre- and postsynaptic elements throughout the neuropil, and the expected peak in pre-to-postsynaptic separation distances consistent with synaptic ultrastructure^54^ (Fig. 7E and S2B).

We next assessed the ability of SEQUIN to quantify synaptic loci in human brain samples, a critical step for translational applications. Samples of formalin fixed human brain from several subjects were sectioned, immunolabeled, imaged, and analyzed (Fig. 7F-I). In samples from each of three subjects a peak in pre-to-postsynaptic separations was identified consistent with synaptic loci (Fig. 7I). These results establish the applicability of SEQUIN to the analysis of synaptic populations with specific afferent origins, molecular and physiological properties, and synaptic loci from human brain samples or those labeled with endogenous fluorescent markers.

### Mesoscale imaging of synaptic microconnectivity

Synapses exhibit population level diversity in density, molecular composition, and physiological properties across mesoscale brain regions (*e.g.*, cortical layers, subregions of hippocampus)^11,55,56^. This variation affects the function of parent networks and may impart unique resilience or vulnerability to disease processes^51^. Capturing such features of synaptic networks requires nanoscopic analysis approaches scalable to mesoscales, ideally to brain regions encompassing functional neural circuits. The natively volumetric imaging and high degree of automation enabled by SEQUIN (Fig. 1D) permits application across such brain regions.

The hippocampal formation is involved in a number of cognitive functions, in particular the storage of spatial information, and is uniquely affected in AD neurodegeneration. To test the ability of SEQUIN to identify mesoscale patterns of synaptic alterations in response to neurodegenerative conditions, we imaged throughout the cortical depth (Fig. S10) and across entire mouse hippocampal sections from control and APP-PS1 animals at super-resolution (Fig. 8A, B). Images consisted of 431 - 583 individual 74 × 74 × 4 μm tiles, and required ~28 hrs to acquire (9.3 sec/1000 μm^3^). Imaging datasets were subsequently batch processed for synaptic localization analysis (a process that required ~2.5 days to complete, with ~90 minutes active time).

**Fig. 8:**
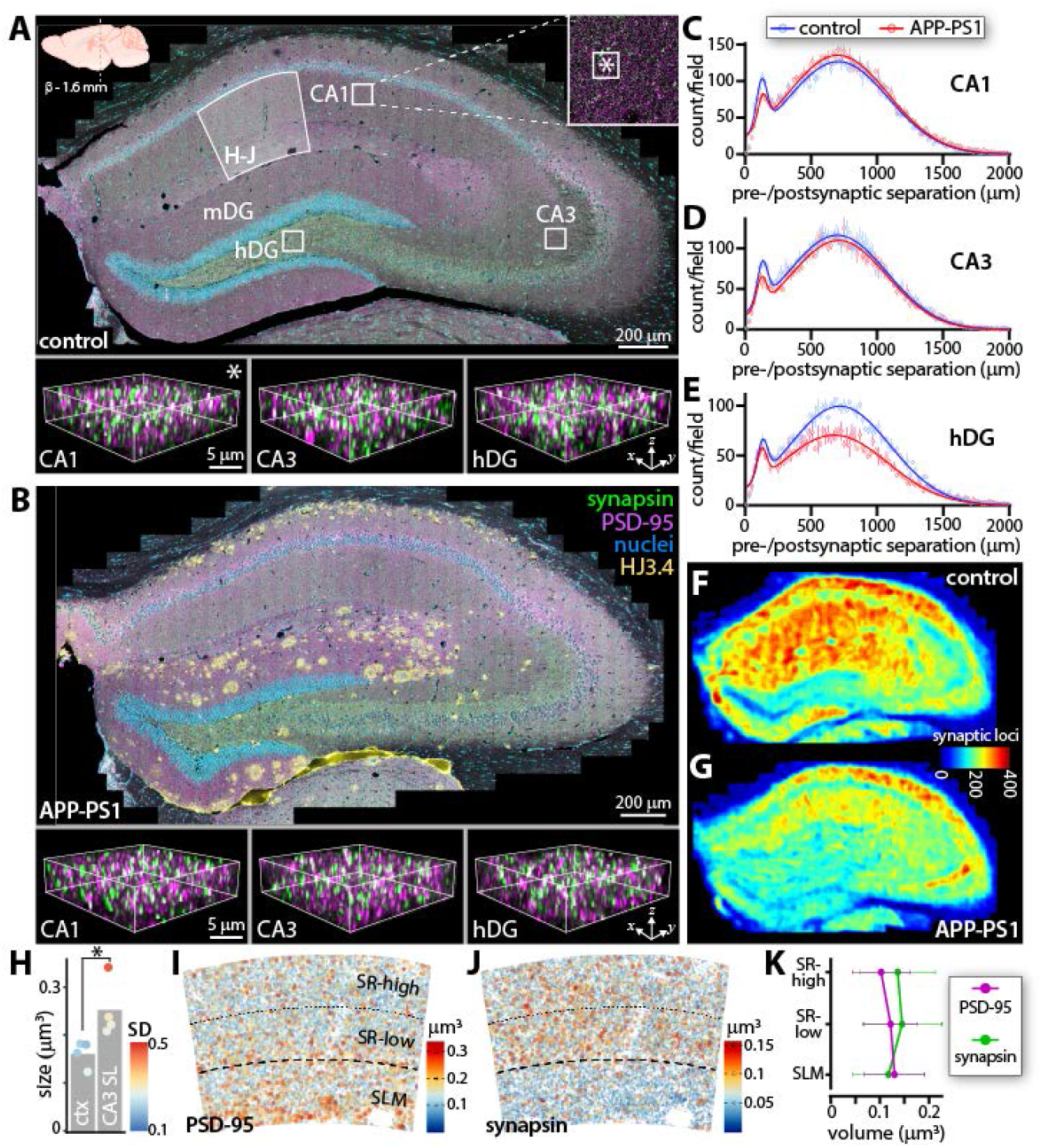
Mesoscale synaptic quantification. (A) Down sampled overview of hippocampal tile scan from control animal. CA1, CA3, and dentate gyrus labeled along with approximate regions for localization analysis in C-E. CA1 boxed region shown in inset, with smaller box (*) indicating size and location of subfield in bottom-left panel. Identically sized subfields from CA3 and dentate gyrus shown in bottom-middle and −right panels, respectively. Region of CA1 subjected to radial pre- and postsynaptic volume comparison in H-J noted. Location of section relative to Bregma indicated in upper-left corner, applicable to A and B. All immunolabels in B apply to A and B. HJ3.4 labels amyloid plaques. (B) Hippocampal tile scan from APP-PS1 animal with subfields highlighted from same regions as in A. (C-E) Frequency distributions of pre-to-postsynaptic puncta separations from boxed regions in A and equivalent regions in B (CA1, CA3, dentate gyrus hilus). Error bars +/−SEM. (F, G) Heatmaps of synaptic density across control and APP-PS1 hippocampi from n = 2 animals each, transformed to a common atlas for averaging and comparison. Color LUT shows number of detected excitatory synaptic loci within a 20 μm XY radius through the scanned depth. (H) Comparison of presynaptic punta size between somatosensory cortex and CA3. Color LUT shows within-animal variability (standard deviation) of puncta sizes from n = 4 animals. (I) Volume of PSD-95 and synapsin (J) puncta in CA1 molecular layer, quantified in (K). * p < 0.05. Abbreviations: hDG, hilus of dentate gyrus; mDG, molecular layer of dentate gyrus; ctx, cortex; SL, stratum lucidum; SD, standard deviation; SR-high, stratum radiatum-superficial; SR-low, stratum radiatum-deep; SLM, stratum lacunosum moleculare.

Inspection of individual images from resulting composite maps revealed resolution of synaptic elements (Fig. 8A, B, lower panels) adequate for quantification of synaptic loci across numerous functionally significant regions of hippocampus (Fig. 8C-E). To better highlight regional variation in synaptic density, and to identify regional susceptibility to synaptic degeneration related to aging and neurodegeneration, we prepared heatmaps of excitatory synaptic density from these datasets. Individual hippocampal heatmaps were then transformed to a common atlas^57,58^ space for averaging and comparison (Fig. 8F, G). Consistent with a report in a non-human primate^59^, detected synaptic loci were denser in CA1 compared to CA3 or the hilus of dentate gyrus. Synapse loss in the AD model was non-uniform: the greatest loss occurred in CA1 and CA3, with relative preservation in the hilus of the dentate gyrus. Correlation of synapse loss with amyloid plaque burden was similarly non-uniform: in the molecular layer of the dentate gyrus, a clear relationship between synapse loss and overall amyloid plaque density was observed, while this relationship was far less apparent in CA1 and CA3, implying multiple spatially-segregated mechanisms of synaptic injury in the hippocampus (see Discussion).

The CA1 molecular layer exhibits radial gradients of synapses with distinct electrophysiological properties that contribute to spatial memory encoding^60–62^. To better conceptualize the significance of regional variation in synaptic structure revealed by SEQUIN for network function, we sought to analyze the sizes of pre- and postsynaptic elements across CA1 laminae. The size of these elements correlates closely with synaptic strength^63–69^, and this nanostructural diversity contributes to hippocampal information storage^70^. Towards this end, we first evaluated the ability of SEQUIN to detect well-described variation in presynaptic bouton size by comparing the large mossy fiber boutons of the CA3 stratum lucidum (SL) with those of somatosensory cortex. Presynaptic elements of the CA3 SL were significantly larger than neocortical puncta on average, though the magnitude of this population-level difference was impacted by the mixture of large with small boutons in CA3^71^, as indicated by the wider distribution of CA3 presynaptic sizes (Fig. 8H).

Turning to the CA1 molecular layer, we found that postsynaptic elements (PSD-95 puncta separated from synapsin puncta by < 200 nm) increased in size with distance from CA1 pyramidal somata (Fig. 8I, K), in agreement with reports using an orthogonal S-R approach^72^. Interestingly, presynaptic puncta (reciprocal to PSD-95 synaptic puncta) in these samples exhibited stable size in proximal CA1 dendritic trees (within striatum radiatum), but were substantially smaller in distal arbors (in stratum lacunosum-moleculare; Fig. 8J, K). This observation suggests a previously unknown size-function relationship in CA1 synapses that may have implications for network behavior and its response to regionally distinct synapse loss. Together, these results establish the utility of SEQUIN as a multiscale synaptic analysis tool that can reveal novel aspects of synaptic distributions, their population-level features, and their differential response to neuropathological processes.

## Discussion

### Synaptic imaging

An expanding collection of innovative histological, imaging, and computational techniques is rapidly enhancing what can be gleaned from light microscopy datasets. While such approaches hold tremendous potential to address critical questions involving nervous system development, function, and malfunction, widespread application is hindered by numerous factors that collectively limit their reach. Synapses offer an illustrative case study: although among the most important of neural structures in health and disease, critical questions regarding their distribution, features, and population level dynamics remain unknown because of their inaccessibility in neural tissue to most imaging methods.

Advanced synapse imaging approaches each present a unique combination of resolution, speed, molecular characterization, and accessibility, among other factors. SEQUIN occupies a particularly useful position within this parameter space. By providing a rapid, easily implemented, artifact resistant^20^ method to quantify synaptic endpoints based on familiar and widely available hardware, SEQUIN should broadly enable the application of synaptic analysis to problems of biological importance. The combination of standard histological processing with exogenous (antibody-based) labeling of synaptic markers furthermore maintains maximum flexibility with parallel experimental determinants, cross-species applications, and archived tissue.

Alternate approaches may be more appropriate for specific questions. EM possesses unparalleled capability to resolve fine synaptic ultrastructure (*e.g.*,^12^). Array tomography may be more suitable when proteomic multiplexing of markers^73^ or their correlation with ultrastructure^16,17^ are top priorities. The exceptional resolution enabled by single molecule localization microscopy approaches (STORM^74^, PALM^75,76^, and related techniques) and stimulated emission depletion (STED)^77^ microscopy can reveal nanoscale features of individual chemical synapses challenging or impossible to detect with image scanning microscopy (*e.g.*, subsynaptic protein nanoclusters^72,78^). STED was also recently shown to be amenable to imaging postsynaptic structures *in vivo* through the use of a knock-in HaloTag fusion to PSD-95 and intracortical injection of a STED-appropriate fluorescent ligand^79^. SEQUIN analysis of natively-fluorescent synaptic fusion proteins (Fig. 7E and S2B) should permit similar *in vivo* investigations without the requirement of injected fluorophores, using conventional or multiphoton excitation. Thus while each technique possesses unique strengths and limitations, the high throughput pipeline and accessibility afforded by SEQUIN permits comparatively straightforward, rapid analysis of synaptic connectivity over mesoscales, while preserving the ability to explore functionally relevant nanoscopic structural and molecular characteristics at the individual synapse and population level.

### Synaptic injury in acute and chronic disease

Synaptic injury and subsequent loss is a shared early feature of many chronic neurodegenerative conditions including Parkinson’s disease, Huntington’s, multiple sclerosis, and AD, among others^2,80,81^. Mutations in >130 human postsynaptic proteins are linked to neurological and psychiatric disease^82^. A reduction in synapse number is also the strongest microstructural correlate of cognitive decline in healthy aging^2,83^. These findings indicate that synapses may be the most vulnerable link in the chain of neurons that constitute functional networks. Understanding this phenomenon is therefore among the most pressing goals of translational neurobiology.

A further common theme in these devastating conditions is the lack of an identifiable trigger that initiates the neurodegenerative cascade. The strong epidemiological link between acute brain trauma and chronic neurodegeneration is, in this regard, likely an important clue. Hazzard ratios for the development of dementia resulting from brain trauma, including but not limited to AD, are 1.29, 2.32, and 4.51 for concussive, moderate, and severe TBI, respectively^9,44,45^. Once thought to be a monophasic injury, TBI is now known to induce chronic neurophysiological alterations and progressive cerebral atrophy in many individuals^84–86^. Recently it has become clear that sufficient traumatic burden on its own leads to a form of dementia—chronic traumatic encephalopathy (CTE)—that shares features with AD yet is a distinct clinicopathological entity^87^.

Prior studies link brain trauma to altered APP and tau metabolism^88–92^. Our findings indicate that diffuse TBI induces synaptic injury and subsequent loss. These observations may be connected: synapse loss may represent a structural-mechanistic link connecting traumatic tau/APP alterations with key dementing neuropathological processes. A direct synaptic susceptibility to biophysical stress has also been proposed^93^. Our findings of spatiotemporally distinct subcellular injury processes in acute brain injury^94^ and across the hippocampus in an AD model suggest that multiple mechanisms may conspire to damage synapses. Clarifying these processes in traumatic and neurodegenerative synaptopathy through the application of SEQUIN is a critical future goal.

Synaptic dystrophy similar to our findings has been reported by EM in contused cortex following TBI in humans^95^. Likewise, perilesional synapse loss has been observed in an animal model of direct cortical contusion through a craniotomy^96,97^. Recently, Vascak and colleagues reported peri-somatic loss of inhibitory synaptic terminals and upstream alterations in axon initial segments in a more clinically relevant model of direct cortical impact lacking macroscale necrotic lesions^98^. Our findings of diffuse, bilateral excitatory synapse loss following TBI in the absence of macroscale lesions further reinforces the developing consensus that traumatic network injury entails far more complex and multifocal circuit disruption than previously appreciated. SEQUIN mesoscale analysis has the potential to both shed further light on these gray matter alterations and connect them with up- and downstream white matter pathways. As all connectional lesions must be addressed to salvage functional circuits, such multiscale approaches will be crucial to the development of future therapeutic interventions. The lack of such perspective may be one reason for the disappointing track record of failed TBI treatments.

### From synapses to connectivity

Synapses connect axons and dendrites into a brain-wide network referred to cartographically as the connectome^99,100^. Large-scale synaptic architecture, the ‘synaptome^56^,’ is both a component of the connectome and may play crucial roles in its development, refinement, systems-level organization, and function. Synapses exhibit exceptional structural, molecular, and functional diversity^56^, with >1400 proteins identified in the PSD^82^ and >400 proteins associated with synaptic vesicles^101^. Proteins expressed in synapses carry the signatures of their pre- and postsynaptic parent neurons^56^. Considering the exceptional number of genes encoding synaptic proteins, and the observation that >70% of genes are expressed in <20% of neurons^57^, capturing this synaptic diversity using mesoscale analysis techniques such as SEQUIN has the potential to reveal crucial features of connectome organization and function.

Zhu and colleagues recently highlighted the power of such an approach using diffraction-limited imaging to capture features of postsynaptic structures canvasing large regions of mouse brain^51^. Synaptic diversity, defined based on imaging features of just two postsynaptic proteins, exhibited remarkable correlation with known structural and functional connectome properties. In computational models, the distribution of these postsynaptic subtypes impacted patterns of postsynaptic activity in response to behaviorally-relevant inputs, implicating synaptic diversity as a means of storing information in and shaping the behavior of neural circuits beyond their hard-wired anatomy. SEQUIN, by virtue of its flexibility, specificity, resolution, and multiscale cataloguing of both pre- and postsynaptic structures would be expected to enhance such efforts to characterize the synaptome, understand its role in coordinating connectome architecture and function, and leverage that knowledge to improve understanding of the neural basis of behavior^56^.

## Materials and Methods

### Animals

All animal experiments were approved by the Washington University Institutional Animal Care and Use Committee and performed in accordance with all relevant guidelines and regulations. Unless otherwise noted, all experiments were performed in 16 wk-old male C57Bl/6 mice (Jackson Labs #000664, Bar Harbor, ME, USA). APPswe/PSEN1dE9^38,39^ (APP-PS1) aged 18 mo were used to quantify synaptic loci in AD. Experimental TBI was generated using the modCHIMERA platform at an impact energy of 2.1 J, as previously described^46^. The mortality rate from injury was 8%. Analysis of endogenous synapse labelling was carried out in Syb2-mRFP^52^ and PSD-95-Venus^53^ double knock-in mice.

### Tissue collection and processing

Tissue for immunofluorescence was collected by anesthetizing animals with isoflurane and transcardially perfusing with ice-cold phosphate buffered saline (PBS) until the liver visibly cleared. Animals were then perfused with ice-cold 4% paraformaldehyde (PFA) in PBS for three minutes. The brain was carefully removed and stored in 4% PFA overnight at 4°C (6 hours for experiments conducted at Ludwig-Maximilians Universität München). The following day the brain was rinsed in PBS and transferred to 30% sucrose in PBS. The brain was left in this solution at 4°C until cutting (at least 48 hrs). Cryostat sections were obtained by placing the brain in a cryomold (Tissue-Tek #4557, Torrance, CA, USA) and surrounding it with a 2:8 mixture of 30% Sucrose in PBS and OCT (Fisher #4585, Hampton, NH, USA). Embedded samples were frozen with dry ice-cooled isopentane, and frozen blocks were stored at −80°C until cryosectioning at 15 μm (Leica CM1950, Buffalo Grove, IL, USA). Sections were collected onto charged glass slides (Azer Scientific, Morgantown, PA, USA. or Globe Scientific, Mahwah, NJ, USA) and kept at room temperature (RT) for at least 30 minutes before storage at −80°C until immunolabeling. Microtome sections were obtained by removing whole brains from 30% sucrose solution, rapidly freezing with crushed dry ice, freezing to the microtome stage with PBS (Microm HM 430, ThermoFisher Scientific, Waltham, MA, USA), and sectioning at 50-150 μm. Sections were stored free-floating in a cryoprotectant solution (30% sucrose and 30% ethylene glycol in PBS) at −20°C until immunolabeling. Human tissue was acquired from 3 subjects with causes of death including nonblast TBI (subjects 1 and 2) and complications of chemotherapy (subject 3). Post-mortem interval for these subjects is unknown.

Tissue collection for electron microscopy (EM) was performed by anesthetizing the animal with isoflurane, transcardially perfusing with 37°C normal Ringer’s solution containing 0.2 mg/mL xylocaine and 20 units/mL heparin for 2 minutes, and then perfusing with 0.15 M cacodylate (pH 7.4) containing 2.5% glutaraldehyde, 2% formaldehyde, and 2 mM calcium chloride at 37°C for 5 minutes. Following perfusion the brain was carefully removed and placed in the same fixative overnight at 4°C. Tissue sections were prepared for EM using the OTO method (osmium and thiocarbohydrazide liganding), as described^102^.

For 15 μm cryostat sections, tissue was rewarmed to RT (~20 min) and isolated with a hydrophobic barrier (Vector Labs ImmEdge #H-4000, Burlingame, CA, USA) that was allowed to dry for ~20 min at RT. For microtome sections, free-floating tissue was placed in 6 well plates with netwell inserts containing PBS. Subsequent steps apply to both cryostat and microtome-cut tissue. Blocking solution, primary and secondary antibody mixtures were centrifuged at 17,000 g for 5 min just prior to use. Tissue was rinsed 3 × 5 min in PBS followed by blocking in 20% normal goat serum (NGS; Vector Labs S-1000, Burlingame, CA, USA) in PBS for 1 hr at RT. Tissue was then incubated overnight at RT with primary antibodies (Table S1) in 10% NGS containing 0.3% Triton X-100 in PBS. The following day sections were rinsed 3 × 5 min in PBS followed by incubation in secondary antibodies (Table S1) in 10% NGS containing 0.3% Triton X-100 in PBS for 4 hr at RT. For nuclear labeling, tissue was rinsed once in PBS followed by incubation in DAPI (1:50,000 in PBS) for 20 min and 3 × 5 min rinsing in PBS. Cryostat sections were briefly rinsed with distilled water prior to coverslipping. Free-floating microtome sections were mounted to charged slides in PBS, dried at RT, briefly rinsed in distilled water, and then coverslipped. Mounting media (~150 μL/slide) was prepared the day of use by mixing Tris-MWL 4-88 (Electron Microscopy Sciences #17977-150, Hatfield, PA) with AF300 (Electron Microscopy Sciences #17977-25) in a 9:1 ratio followed by vortexing and bench top centrifugation to remove bubbles. High precision 1.5H coverglass was used for all experiments (Marienfeld # 0107242). Coverglass was thoroughly cleaned with ethanol and air dried prior to use. Slides were allowed to cure protected from light at RT for at least one day before the short edges of the coverglass only were sealed with CoverGrip (Biotium #23005, Fremont, CA, USA).

For sections thicker than 50 μm, primary antibody incubation was extended to 3 days (with sodium azide at 0.2%) and secondary antibody incubation to 12 hrs. NGS was replaced with 4% bovine serum albumin when primary antibodies raised in goat were used. Anti-goat secondaries were rinsed extensively before the addition of secondaries raised in goat.

### Image acquisition

Images were acquired on a Zeiss LSM 880 microscopy with AiryScan detector (Zeiss, Oberkochen, Germany). Either 63x 1.4NA (primary) or 100x 1.46NA (secondary) oil immersion objectives were used for all experiments. Images were acquired using an optical magnification of at least 1.8x. XY pixel size was optimized per Zeiss Zen software (43 nm pixel size). Z pixel dimension was set to modestly oversample (120 nm Z-step; 30% oversampling) or to optimal (180 nm Z-step; for hippocampal tile scans). Laser power and Z-ramp (when necessary) for each channel was set independently to equalize signal at the top and bottom of the imaging stack and to occupy the first ~1/3 of the detector dynamic range. Scan speed was set to maximum, scan averages to 1, gain to 800, and digital gain to 1. Confirmation of AiryScan detector alignment was performed before image acquisition and was rechecked with every new slide. Hippocampal mesoscale imaging was conducted at the center of 50 μm microtome sections. The convex hull function in Zeiss Zen was used to outline the borders of the hippocampus defining the region to be imaged prior to unsupervised, automated image acquisition. Cortical mesoscans (synapsin or vGluT2 and PSD-95) were acquired similarly. During each imaging session, a within-tissue, within-antigen, within-experiment channel alignment control was also imaged. This consisted of an experimental brain section labeled during each experiment against synapsin and a 1:1 mix of secondary antibodies bearing all spectrally-distinct fluorophores used in the experiment. All experimental image acquisition involving group comparisons was conducted in a fully blinded fashion.

### Image pre-processing

Following acquisition, images were 3D Airyscan processed in Zeiss Zen Black. For TBI experiments, the Wiener parameter (10^-f^) was set with an f value (the Airyscan filter strength) for the entire experiment equivalent to the lowest automatically-determined value in Zen Black. This and subsequent imaging experiments revealed that the automatically-determined f value was generally very close to 6, and that departures within the range of automatically-determined values had no substantial impact on synaptic quantification. We therefore used a fixed f value/Airyscan filter strength of 6 for all other experiments. Following Airyscan processing, the channel alignment images were inspected to determine the magnitude of lateral and axial chromatic aberration for that imaging session. A manual correction was applied in Zen Black to eliminate misalignment, and this correction was propagated to all experimental images.

### pre- and postsynaptic puncta and synaptic loci analysis

Pre- and postsynaptic puncta detection: Puncta detection was performed using Imaris software (Bitplane, Zurich, Switzerland), version 9. The spots detection function was applied to each channel. Spot size was adjusted to maximize detection of visually evident puncta (XY size of 0.2 μm for rabbit anti-PSD-95 labeling, and 0.09 μm for goat + rabbit anti-PSD-95 labeling) with integrated background subtraction. A Z size of 0.6 μm (rabbit anti-PSD-95) or 0.27 μm (goat + rabbit anti-PSD-95) was used for spots parameter extraction. Following spots identification, a 0.1 μm XY, 0.3 μm Z guard was applied to eliminate spots clipped by the image border from further analysis (to ensure a maximally inclusive dataset for subsequent analysis, no further filters were applied). Bitplane Batch Coordinator was utilized for mesoscale imaging to automate these processes. Signal resulting from autofluorescence was eliminated using the region growing feature in Imaris to eliminate spots larger than 0.33 μm^3^ in the red channel. Region growing was also applied using an automatically-determined absolute intensity threshold to determine pre- and postsynaptic puncta volumes in hippocampus and cortex (Bregma −1.6 mm). All such measurements were internally controlled. The XYZ position of the centroid of every spot, along with extracted parameters, was output to spreadsheets for further processing. In some cases (to assess purely random puncta associations; Fig. 3G, I and Fig. 4B, C) the postsynaptic coordinates were reflected in the Y dimension prior to synaptic loci detection.

Synaptic loci identification: Nearest neighbor analysis was performed using custom scripts in MATLAB (Mathworks, Natick, MA, USA) to analyze the XYZ centroid position and mean intensity of all puncta. The SEQUIN code first applied a user defined intensity range filter to select the desired spots for nearest neighbor calculation. The 20% most intense puncta for both pre- and postsynaptic channels were used for all experiments except those in Fig. 4, S6, and S7 in which multiple intensity ranges were assessed. The SEQUIN code then calculated the 3-dimensional Euclidean distance (*d*, Eq. 1) between each postsynaptic punctum and its nearest presynaptic neighbor (run with a presynaptic reference for Fig. S7).

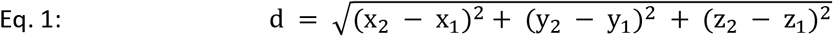

Linear DNA origami nanorulers consisting of two Alexa Fluor 568 fluorophore markers each 70 nm from a central Alexa Fluor 488 fluorophore marker (GATTAquant SIM 140YBY, Hiltpoltstein, Germany) were analyzed using SEQUIN localization analysis. Images were initially spatially oversampled (pixel size 9 × 9 × 20 nm XYZ), then downsampled to standard SEQUIN pixel size for localization analysis.

Synaptic loci quantification: A cut-off of 200 nm pre-to-postsynaptic separation was used to quantify synaptic loci for all experiments outside of Fig. 4 (400 nm), in which a nearest neighbor frequency distribution resulting from reflected postsynaptic data was first subtracted from the original frequency distribution of pre-to-postsynaptic separations (see Results; Fig. 4B, C). To quantify synaptic loci via Gaussian unmixing, the Nonlinear Least Squares function in R (The R Foundation, Vienna, Austria) was used to derive mean, amplitude (*A*), and standard deviation (σ) of the underlying Gaussian distributions. Fits were visually inspected, and synaptic density calculated from the Gaussian distribution representing the early peak in the pre-to-postsynaptic separation frequency distribution (Eq. 2; see Fig. S6).

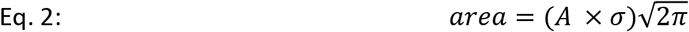

Pathological synapse loss: Quantification of diffuse synapse loss after experimental TBI or in APP-PS1 mice was performed in layer 1 cortex at Bregma −1.6 mm (primary somatosensory cortex). For analysis in APP-PS1 mice, 16 week old male C57Bl/6 mice were used controls. Strict blinding was maintained during imaging and image analysis. Two images were collected from both the left and right sides of the brain, for a total of four images per animal. The number of detected synaptic loci were averaged on a per animal basis for groupwise comparison. Quantification of synaptic density as a function of distance from amyloid plaques was carried out in 50 μm microtome sections from APP-PS1 mice labeled against PSD-95, synapsin, and amyloid plaques (HJ3.4; Table S1). Following imaging, the XY center position of each plaque was manually marked and used for subsequent plaque position-relative quantification of synaptic loci within ~7 μm-wide rings centered on the plaque core.

Spatial statistics: Spatial statistics of pre- and postsynaptic puncta and synaptic loci were analyzed using the Spatstat package^103^ in R. Ripley’s K function^32^ were applied using the *Kest* (univariate; synapsin or PSD-95) and *Kcross* (bivariate; synapsin and PSD-95) functions. Nearest neighbor orientations of synaptic loci were analyzed using the *nnorient* function, and the Fry plot was calculated using the *fryplot* function.

Heatmaps of synaptic density: To create heatmaps of identified hippocampal synaptic loci, background subtraction was first applied by eliminating pre- and postsynaptic puncta < 2x as intense as the auto-determined (Imaris) diffuse background level in white matter, followed by SEQUIN identification of synaptic loci. The XY positions of resulting synaptic loci (defined as the centroid of the postsynaptic punctum) were represented as 20 μm circles with the same center and of gray value 1. Overlapping circles were summed to derive raw heatmaps wherein each pixel represents the density of synaptic loci within a circle of radius 20 μm. Raw heatmaps were transformed to reference atlas^57,58^ space in ImageJ using the Landmark Correspondences tool (20 landmarks placed on the reference atlas and each raw heatmap). A moving least squares affine transformation was then performed. vGluT2 heatmaps were generated using the ggplot2 and hexbin packages in R (geom_hex) using a binwidth of 20 μm.

### Electron microscopy

Electron micrographs with 10 nm resolution were acquired on a Zeiss Merlin FE-SEM. For measurement of distances between pre- and postsynaptic marker equivalents, synapses were first identified by the presence of a clear presynaptic vesicle pool and post-synaptic electron density. Both the vesicle reserve pool and the PSD were manually traced in ImageJ and the XY center position of these traces were used to calculate the trans-synaptic Euclidean distance.

Quantification of synaptic density after experimental TBI was performed in a fully blinded fashion on electron micrographs with 7 nm resolution. Systematic random sampling was performed by first organizing the raw images onto an XY grid of non-overlapping imaging fields. A square dissector was then overlaid on each individual image containing two inclusion and two exclusion borders. Synapses were identified as above. Synapses were counted if the vesicle pool was fully inside the dissector or touching the inclusion border. Synapses were excluded if the vesicle pool touched the dissector’s exclusion border.

### Statistics

Parametric statistics were used for all tests except where group numbers were n < 5, a test of normality (Shapiro-Wilk test), or visual inspection of the data indicated a non-normal distribution, as noted below. All tests were two sided except the EM analysis of synaptic density after experimental TBI, where an exclusive, pre-specified directionality based on SEQUIN analysis was of interest. α < 0.05 was used to determine statistical significance for all comparisons. Analysis of nanorulers (Fig 2F), synaptic density by EM post-TBI, and cortical synaptic loci in APP-PS1 mice (Fig 5D-F) involved n < 5 subjects. The Mann-Whitney U-test was therefore utilized. SEQUIN analysis of synaptic loci after TBI (Fig 6F) and analysis of cortical thickness (Fig. S8B, C) was carried out with a one-way ANOVA with Holm-Sidak post-test. The Mann-Whitney U-test was also used for groupwise comparisons of cortical vs. hippocampal presynaptic puncta volumes. Elsewhere data are summarized as mean ± SEM. All groupwise comparisons were assessed in a blinded fashion.

## Supporting information

SEQUIN supplemental information

## Acknowledgements

The authors wish to thank Drs. Jens Rettig, Haining Zhong, and Hao Jiang for providing the synaptobrevin and PSD-95 endogenously-labeled mouse lines, Dr. James Fitzpatrick and Matthew Curtis for critical discussions of imaging approaches, Drs. Krikor Dikranian and Joseph Ojo for discussion of electron microscopic images, and Dr. James Fitzpatrick for critical appraisal of the manuscript.

## Author Contributions

ADS participated in project design and co-wrote the MATLAB scripts, performed the experiments and data analysis, and contributed to manuscript preparation. MG co-wrote the synapse analysis code and participated in data analysis. SJR, MHS, SHK, and TM performed histological processing, imaging (SJR), and participated in data analysis. CH and MK independently replicated the core SEQUIN analysis. DLB participated in manuscript preparation and project design. TTK conceived of the project and was primarily responsible for project direction, performed data analysis, wrote the manuscript (with ADS), and oversaw all aspects of the study.

