## Supplementary material for "SEQUIN multiscale imaging of mammalian central synapses reveals loss of synaptic microconnectivity resulting from diffuse traumatic brain injury": SEQUIN supplemental information

### Supplementary Information

#### Supplemental Methods

Refractive index matching

Images of entire sections to demonstrate RI matching were generated using an Epson (Perfection V33) flatbed scanner and a reference grid (2 mm x 2 mm). PBS and Mowiol 4-88 mounted sections were scanned using identical parameters. Plots of intensity by depth were generated in ImageJ using the 'Plot Profile' function on a representative orthogonal projection.

SEQUIN independent replication

All sample processing, imaging, and analysis was conducted in an independent laboratory at a separate institution (Ludwig-Maximilians Universität München, Munich, Germany). Tissue samples for analysis were obtained from offspring of C57Bl/6J-Biozzi and Thy1 GFP line-M[104] mouse line crosses. Anti-synapsin and anti-PSD-95 (raised in rabbit) antibodies were per Table S1. Secondary antibodies were conjugated to Alexafluor 555 and 647 (Invitrogen A21435 and A21244, Carlsbad, CA, USA). Images were obtained on a Zeiss LSM800 microscope with Airyscan detector. Processing was performed using Imaris version 8 (Bitplane, Zurich, Switzerland) and MATLAB (Mathworks, Natick, MA, USA) using identical processes and scripts.

Intensity invariance of puncta

To evaluate intensity ranges both dimmer and brighter than typically utilized, a fixed region was repeatedly imaged at varying laser power from low to high to minimize photobleaching. To avoid any quantitative confounds from decreased signal as a result of potential photobleaching, the final analysis was based upon the mean intensity of the image rather than the laser power used to acquire it.

Cortical thickness measurements

Measurement of cortical thickness was performed on Nissl stained modCHIMERA tissue collected at 7 and 30 days post-injury. Images were acquired on a Zeiss Axioscan automated slide scanner with a 5x objective from sections located at Bregma -1.6 mm. Following image export, Image J was utilized to measure the thickness of the entire cortex as well as layer 1. All measurements were blinded. For the measurement of total cortical thickness, a measurement line was drawn parallel to the central sulcus from the superior-most extent of the corpus callosum (the cingulum bundle) to the cortical surface (Fig. S8a; green line). For the measurement of layer one cortex thickness, a measurement line overlapping the prior line was drawn from the deep border of layer one cortex to the cortical surface (Fig. S8a; red line).

**Supplemental Table**

| | Target | Vendor | Product # | Host | Concentration | Fluorophore |
|---|---|---|---|---|---|---|
| Primary antibodies | PSD-95 | Invitrogen | 51-6900 | Rabbit | 1:200 | N.A. |
| | PSD-95 | Abcam | 12093 | Goat | 1:200 | N.A. |
| | Synapsin1/2 | Synaptic Systems | 106004 | Guinea Pig | 1:500 | N.A. |
| | Homer 1 | Synaptic Systems | 160006 | Chicken | 1:500 | N.A. |
| | vGluT2 | Synaptic Systems | 135404 | Guinea Pig | 1:500 | N.A. |
| | Gephyrin | Synaptic Systems | 147011 | Mouse | 1:500 | N.A. |
| | HJ3.4 | Holtzman lab[a] | HI3.4 | Mouse | 1:1000 | N.A. |
| Secondary antibodies | Rabbit IgG | Invitrogen | A11037 | Goat | 1:200 | Alexa 594 |
| | Guinea Pig IgG | Invitrogen | A11073 | Goat | 1:200 | Alexa 488 |
| | Goat IgG | Invitrogen | A11058 | Donkey | 1:200 | Alexa 594 |

**Supplemental Table 1: Primary and Secondary antibodies used for experiments.**

[a]HJ3.4 was purchased from the laboratory of Dr. David Holtzman, Washington University, St. Louis, MO

**Supplemental figures**

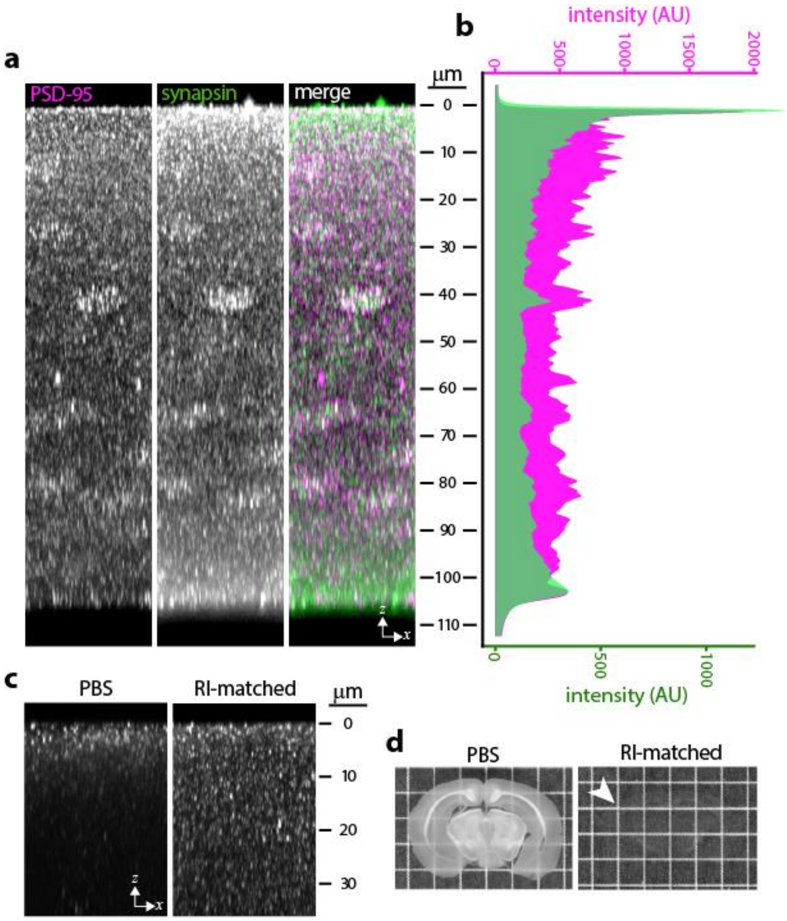


**Fig. S1: Antibody penetration and refractive index matching**

**(a)** 150 µm-thick murine brain sections were immunolabeled in solution, resulting in labeling throughout section thickness; quantified in **(b)**. **(c)** Signal lost in <10 µm of cut surface without refractive index (RI) matching, which effectively clears brain sections **(d**; arrowhead highlights edge of brain section; grid size 2 mm**)**. Data representative of n = 3 animals (a-c) and >10 animals (d).

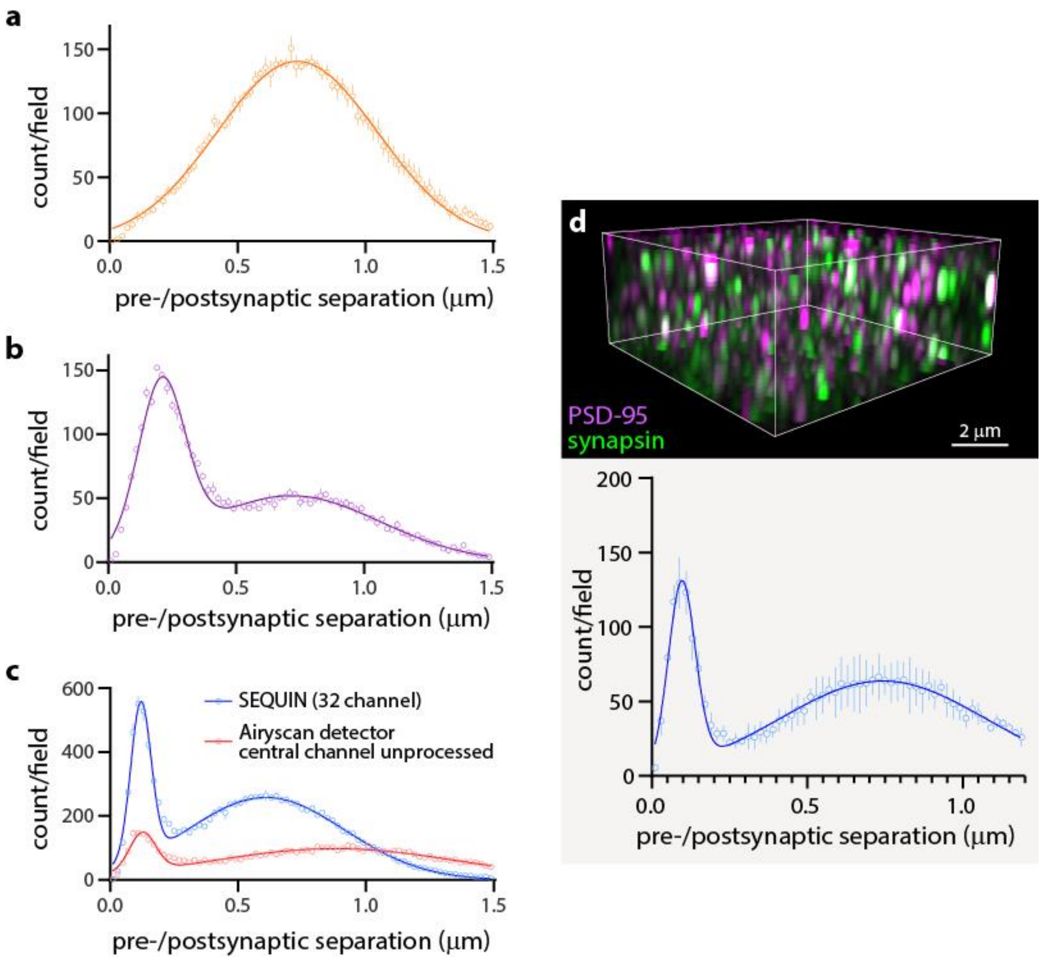


**Fig. S2: Spectral cross-talk, antibody-related and image processing artifact controls, and independent laboratory replication**

**(a)** When the primary antibody for PSD-95 is omitted during immunolabeling, no early peak in pre-to-postsynaptic separations is identified, eliminating the possibility of spectral cross-talk resulting in co-associated pre- and postsynaptic puncta detections. **(b)** Using genetically-encoded labels only (no immunolabeling; see “Analysis of distinct synaptic subsets” section of Results), a bimodal distribution of pre-to-postsynaptic separations can be identified. Thus this pattern is not dependent on antibody-related artifacts. **(c)** Although inefficient due to poor light gathering capabilities and substantially noisier requiring greater intensity filtering, the central channel of the compound eye detector alone yields a bimodal pre-to-postsynaptic separation frequency distribution without requiring post-acquisition image processing. **(d)** SEQUIN imaging and analysis of mouse cortex performed by an independent laboratory following the protocols outlined in Fig. 1. Error bars +/-SEM from n = 3 animals (a-c) and 4 animals (d).

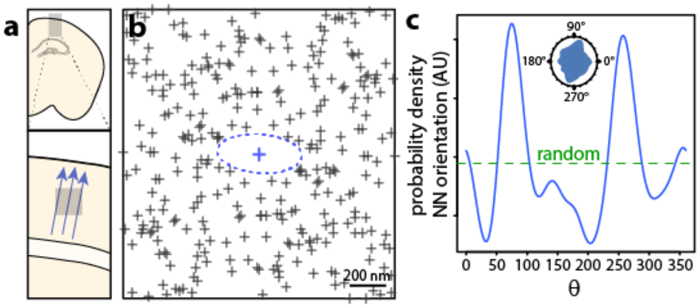


**Fig. S3 Radial orientation of synaptic loci in murine cortex**

**(a)** Schematic of pyramidal neuron dendritic radial orientation in murine cortex. **(b)** Fry analysis of neighboring synaptic loci. Central region of exclusion (blue dotted outline) represents distance between each synaptic locus in the field and its nearest neighbor in all directions. Elongated axis perpendicular to expected radial orientation of synaptic loci is consistent with more distant spacing along this orientation, and with closer nearest neighbor spacing radially along pyramidal dendrites. **(c)** Probability density of nearest neighbor orientation as a function of angle. Note higher probability of orientations aligned radially with cortical anatomy (see rose plot inset). Examples representative of n = 3 animals.

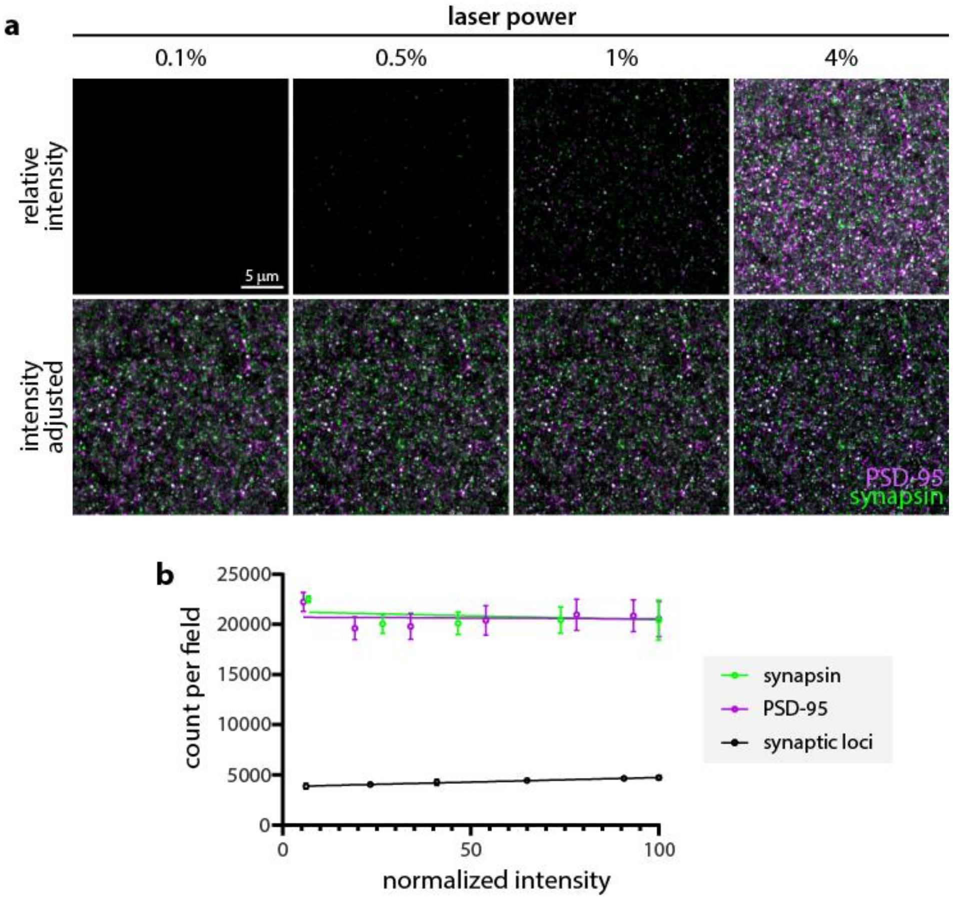


**Fig. S4: Intensity invariance of synaptic puncta detection**

**(a)** Airyscan images of pre- and postsynaptic puncta at varying laser powers before (upper panels) and after (lower panels) manual post-acquisition intensity adjustments. **(b)** Puncta and synaptic loci detection on unadjusted images of varying intensity (resulting from laser power adjustments during acquisition). Note stability of detection across large intensity range. Error bars +/-SEM from n = 14 images from 2 animals.

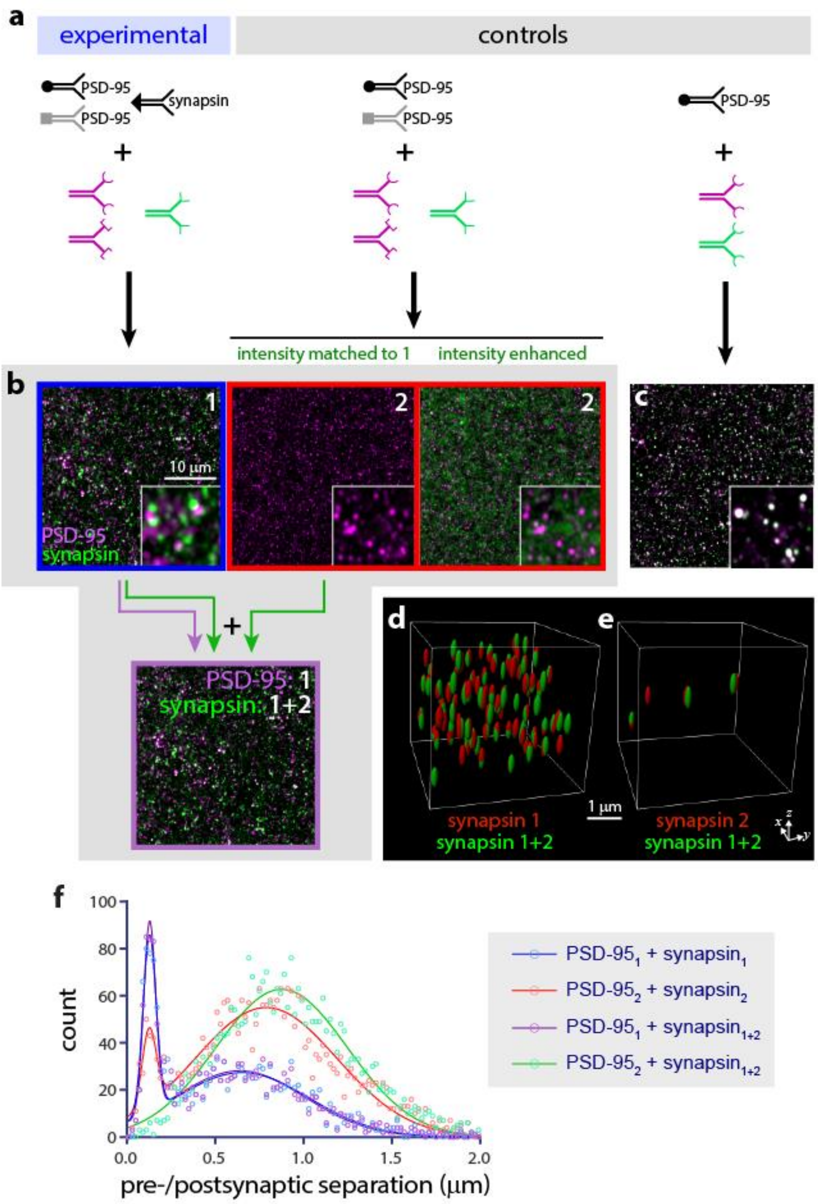


**g**

| | total synapsin puncta detected | synapsin puncta co-located with 1+2 |
|---|---|---|
| 1 | 36,414 | 30,453 (83.6%) |
| 2 | 47,474 | 1,860 (3.9%) |
| 1+2 | 37,020 | identity |

**Fig. S5: Synaptic antibody cocktail controls**

**(a)** Schematic depicting sequential immunolabeling for experimental tissue and controls. **(b)** Labeling against PSD-95 (magenta) and synapsin (green) in experimental tissue (1, blue outline). In controls lacking synapsin primary antibody (2), no synapsin labeling is observed (left red outlined panel) unless intensity is enhanced post-acquisition revealing non-punctate green background (right red outlined panel). To evaluate the effect of this background on puncta and synaptic loci detection, the green channels from images 1 and 2 were combined with the PSD-95 signal from experimental tissue (b, lower panel). Panel outline colors correspond to trend fit lines in (f). Insets (b-c) show subregions at higher magnification to highlight staining pattern; scale bar applies to (b-c). **(c)** Appearance of PSD-95 puncta intentionally cross-labeled with two spectrally distinct fluorophores (magenta + green = white). This pattern does not resemble that of experimental tissue or control lacking synapsin primary. **(d)** Synapsin puncta from experimental tissue (1) co-localized within 2 pixels of synapsin puncta detected from combined (1+2) image, revealing high degree identity. **(e)** Puncta in green channel detected from control tissue (2) minimally co-localize with synapsin puncta detected from combined (1+2) image (quantified in g). Only co-localized puncta (1 or 2 vs. 1+2) are shown. **(f)** Frequency distributions from datasets in (b). Note that although an early peak can be detected in the control lacking synapsin with this antibody combination (red line), adding this background to the experimental signal, as expected from the K-means clustering puncta detection method, results in almost no alteration in brighter experimental synaptic loci detection (compare blue and purple lines). Accordingly, PSD puncta detected from control tissue (2) fail to co-localize with synapsin puncta detected from the combined (1+2) image at separation distances consistent with synapses (no early peak in green line). **(g)** Quantification of above images confirms minimal detection of background puncta in the presence of experimental signal, leading to a maximum false positive rate of 5.9%, and a maximum false negative rate of 19.2% (average of n = 3 cohorts). True false positive and false negative rates are expected to be far lower as the majority of potential noise puncta erroneously detected or experimental puncta erroneously missed will not pair at synaptic separation distances (21.2% of experimental puncta pair at separation distances consistent with synapses in these images). Example data representative of n = 3 cohorts.

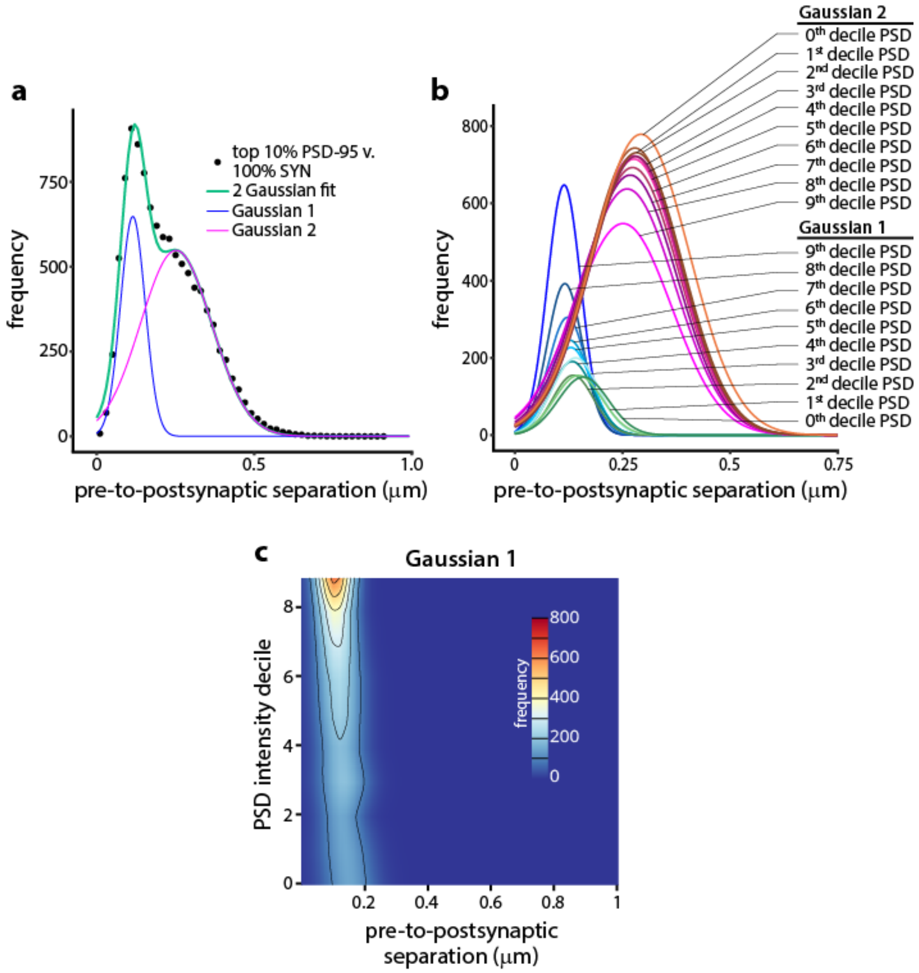


**Fig. S6: 2-Gaussian analysis of SEQUIN frequency distribution**

**(a)** Example of 2-Gaussian fit to SEQUIN frequency distribution and underlying distributions. **(b)** First and second Gaussian components of 2-Gaussian fit for all PSD-95 puncta intensity deciles paired with 100% of synapsin puncta. Gaussian 1 reflects pairings at separation distances ultrastructurally consistent with synapses. Gaussian 2 reflects random pairings. **(c)** Heatmap of pre-to-postsynaptic puncta separations as a function of PSD-95 puncta intensity decile for Gaussian 1. Quantification shown in Fig. 4c. n = 5 animals.

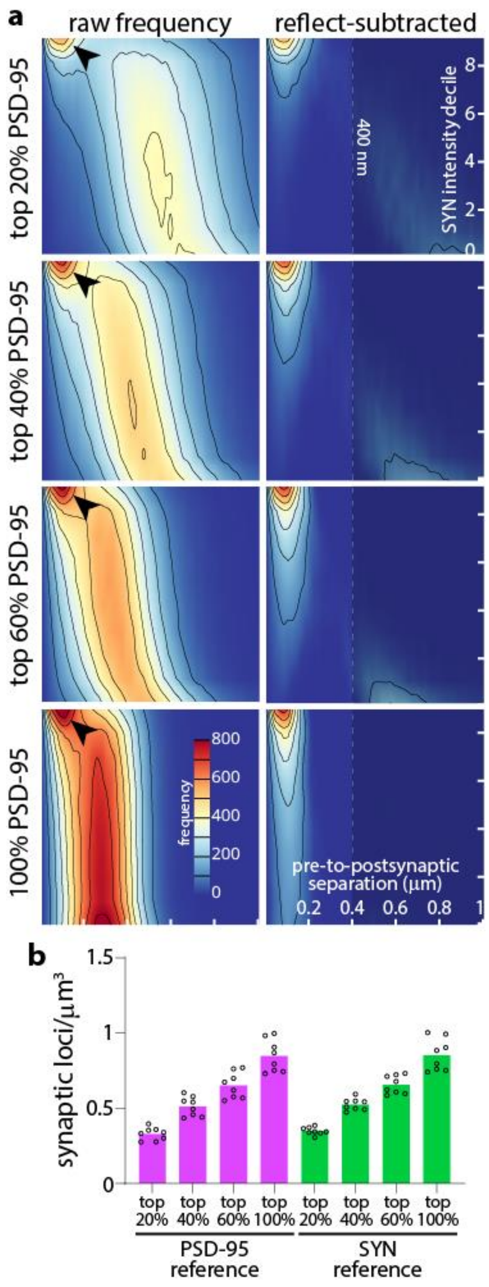


**Fig. S7: Synaptic density quantification with synapsin reference**

**(a)** Heatmaps of pre-to-postsynaptic puncta separations as a function of synapsin puncta intensity decile paired with increasingly inclusive PSD-95 puncta intensity bins. Arrowheads indicate peak of separations consistent with synapses (left panels), which are isolated by subtraction of reflected frequency distributions (right panels). **(b)** Quantification of synaptic loci (closer than 400 nm in reflect-subtracted frequency distributions) yields estimates that closely parallel those observed with PSD-95 reference quantifications. n = 8 animals. Abbreviations: SYN, synapsin.

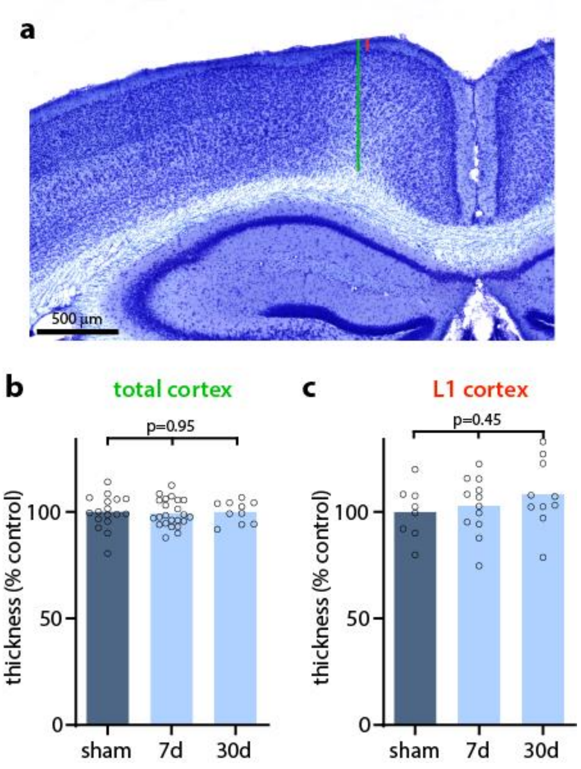


**Fig. S8: Lack of cortical or layer-specific atrophy after modCHIMERA TBI**

**(a)** Brain section demonstrating method of measuring total cortical (green line) and layer 1 (red line) thickness superior to cingulum bundle. There was no difference in thickness of total cortex generally **(b)** or layer 1 cortex specifically **(c)** at 7 or 30 days post injury compared to controls. n ≥ 8 animals/group.

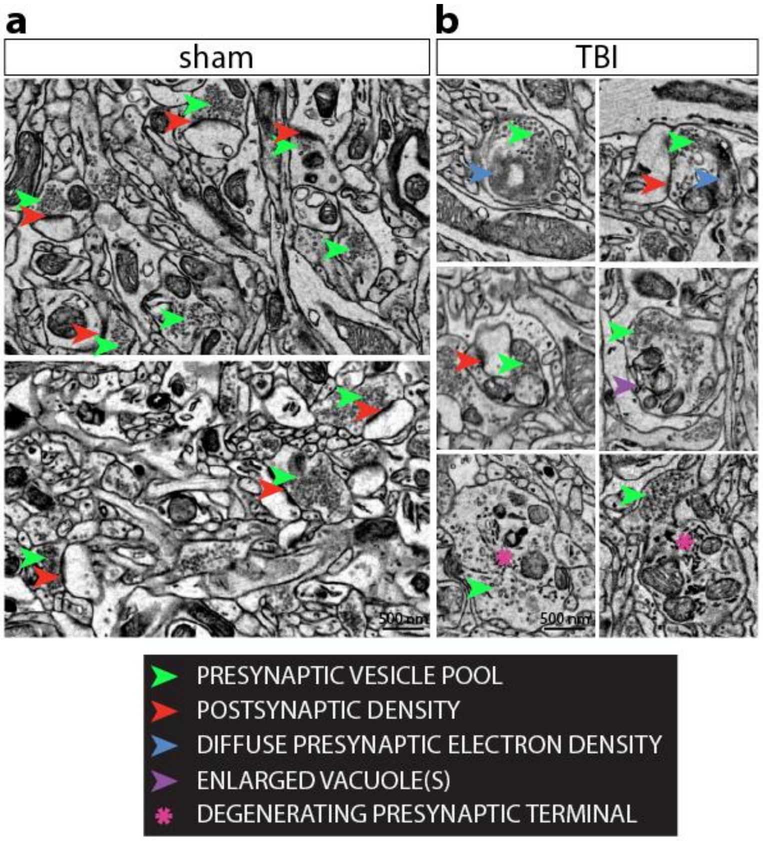


**Fig. S9: Synaptic ultrastructure after modCHIMERA TBI**

**(a)** Further examples of intact synapses from control animals. **(b)** Additional examples of dystrophic synapses identified 7 days after diffuse TBI induced with modCHIMERA. Scale bars in (a) and (b) apply to all panels. n = 3 animals/group.

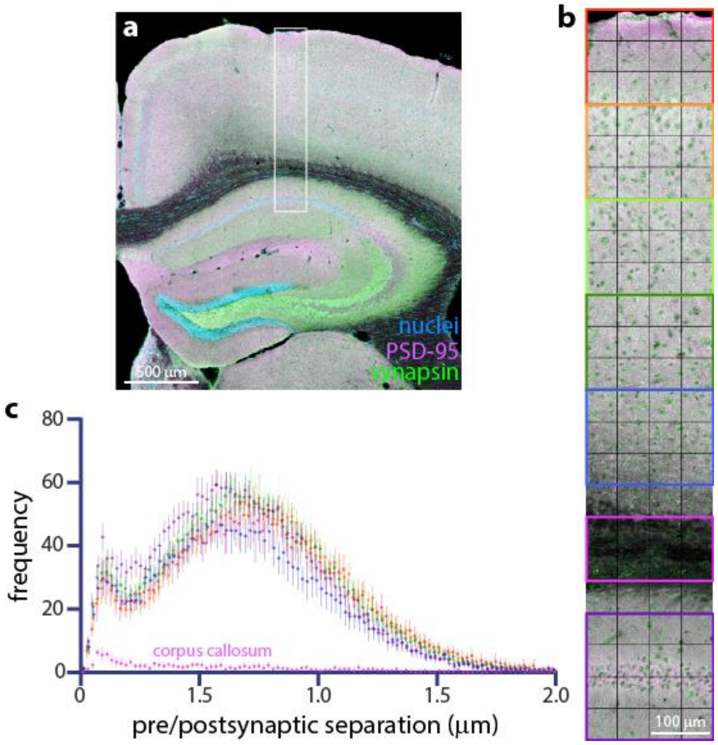


**Fig. S10: Quantification of synaptic loci across several layers of neocortex**

**(a)** Low power image of mouse brain section with region used for quantification boxed. **(b)** Higher power image of cortical depth tile scan with individual subregions quantified in C boxed. Colors correspond to points in (c). **(c)** Frequency distribution of pre-to-postsynaptic puncta separation reveals applicability of SEQUIN to multiple layers of neocortex and hippocampus. Error bars +/-SEM from images within subregion. Data representative of n = 3 animals.
